# Synaptic-state-dependent plasticity routes learning signals in recurrent circuits

**DOI:** 10.64898/2026.06.10.731456

**Authors:** Denis Turcu, Jonathan Cornford, Sven Dorkenwald, Stefan Mihalas

## Abstract

Synapses differ in molecular and ultrastructural state, but the computational role of this heterogeneity remains unclear and computational models do not account for it. Local Hebbian-like plasticity is well supported experimentally, whereas learning complex behaviors most likely requires non-local credit-assignment signals. Recent cortical electron microscopy data suggest that strong and weak synapses occupy distinct structural states, with large synapses often containing a spine apparatus that can alter calcium dynamics and plasticity. Here, we assigned different computational roles to different morphological synaptic types: we asked how networks with state-dependent plasticity operate and whether the mechanisms and structural features they develop are consistent with biological observations. We built a plasticity-switching recurrent neural network (psRNN) in which synaptic state is modeled by recurrent-weight thresholds. Synapses in the low-weight state use a local Hebbian-like rule, whereas synapses in the high-weight state receive an idealized non-local credit-assignment signal implemented with back-propagation through time. psRNNs learned cognitive tasks with temporal and working-memory demands in fewer trials than fully backpropagation-trained RNNs, despite routing BP-based updates to fewer recurrent weights. The advantage arose from three interacting mechanisms: 1) Hebbian-like updates exposed the credit-assignment component to several nearby parameter configurations, 2) the network dynamically built a task-relevant recurrent initialization, and 3) synapses switched into the credit-assignment subset before Hebbian growth destabilized learning. Structurally, trained psRNNs produced more anti-symmetric weights, lower effective rank, and stronger feedforward structure than control RNNs. Our results shed light on the computational role of heterogeneous synapses, which may regulate distinct types of learning signals, and we generate falsifiable predictions for connectomic analyses of recurrent cortical circuits.

## 1 Introduction

Synapses are not interchangeable learning sites. Their size, molecular composition, and organelle content vary across cortical circuits, and these differences can change the plasticity rules available to individual synapses^1–4^. This heterogeneity creates an opportunity for computation: instead of applying one learning rule uniformly across a circuit, the brain could route different learning signals to synapses in different states.

In the past decades, appreciation for the various computational mechanisms of the brain has been growing considerably. For example, large bodies of work have focused on the heterogeneity of cell types and their distinct functional roles; computational models followed this trend, providing key insights into how neural circuits compute distributively^5–12^. In recent years, new observations more clearly pointed out distinct synaptic states^13^ likely undergoing different plasticity mechanisms^3^. However, computational models have yet to explore the functional, computational, and structural consequences of synaptic heterogeneity.

Recent cortical connectomic data suggest a possible substrate for selective routing of learning signals. In the MICrONS phase 1 connectome, cortical synapses between pyramidal neurons are well described by two structural states: small synapses that typically lack a spine apparatus and larger synapses that often contain one (Fig. 1a)^13^. The spine apparatus, a smooth endoplasmic reticulum-derived organelle, acts as an internal calcium reservoir and can influence synaptic plasticity dynamics^1–3^. Plasticity pathways that depend on internal calcium stores, including *IP* 3-dependent *Ca*^2+^ release, can therefore differ between synapses with and without endoplasmic reticulum^4^. These observations raise a concrete computational hypothesis: synaptic structural state may regulate which synapses use local, relatively simpler, plasticity and which synapses receive non-local, relatively complex, credit assignment signals.

**Figure 1:**
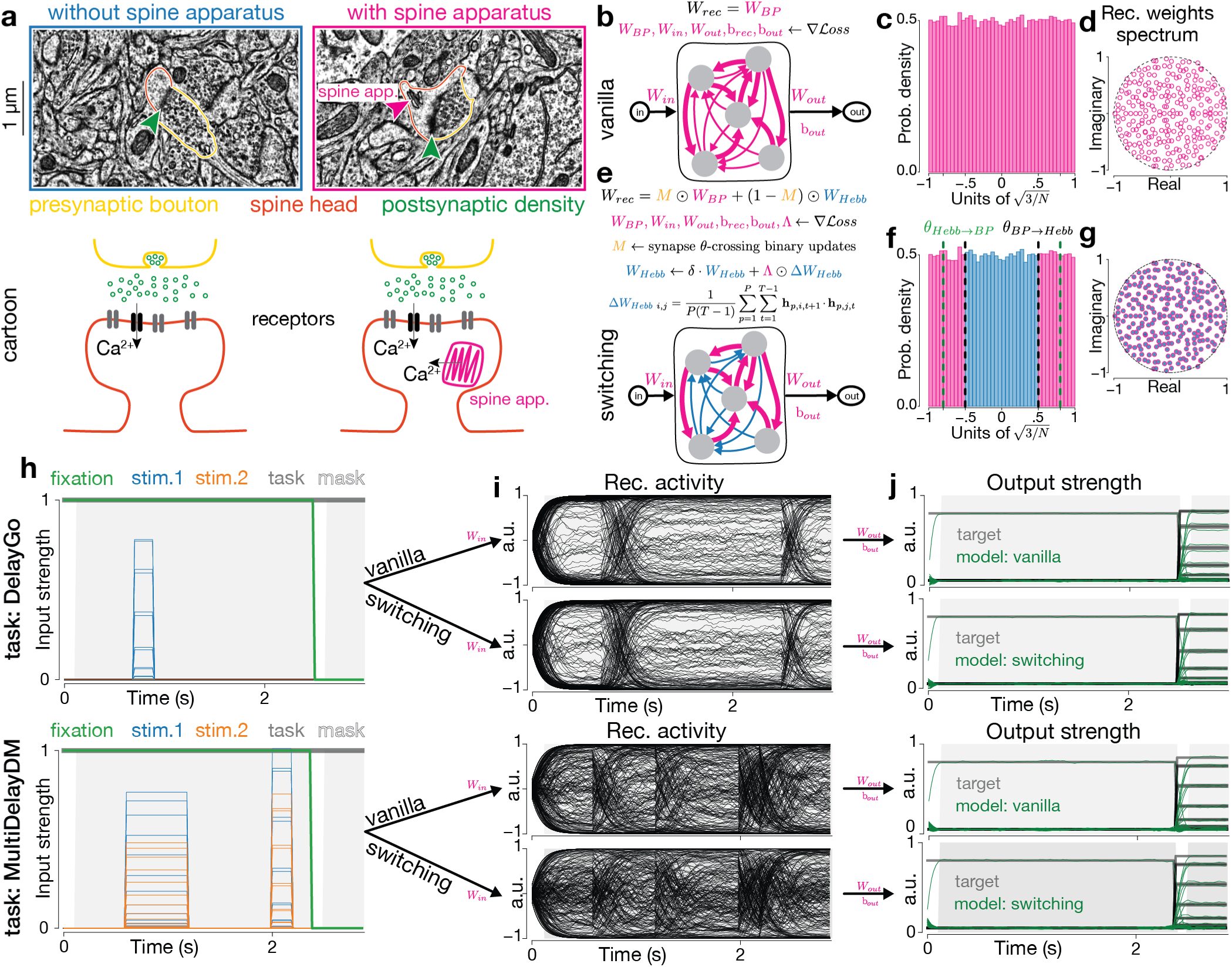
Synaptic states motivate a plasticity-switching recurrent network. **a** Transmission electron microscopy images of example synapses with and without a spine apparatus (top). Cartoon illustration of the two synaptic states (bottom). The presynaptic boutons and spine heads are contoured in yellow and orange, respectively. The postsynaptic density is marked by green arrow head. The spine apparatus is marked by pink arrow head. The cartoon includes a schematic of receptors, with a highlighted *Ca*^2+^ channel. The spine apparatus (pink) provides an additional, internal *Ca*^2+^ source. **b** Control RNN architecture (vanilla). All parameters (in pink) are trained using backpropagation through time (BP). **c** Recurrent weights distribution at initialization. Recurrent weights are sampled from a uniform distribution with a scale that sets the spectral radius at initialization to ∼ 1. **d** Spectrum of the recurrent weight matrix at initialization. Dashed line indicates the unit circle. **e** Switching RNN architecture. Parameters in pink receive BP updates, parameters in blue are updated using a local Hebbian-like rule, and the switching mask in yellow is updated based on synaptic strength throughout training. **f** Recurrent weights at initialization same as vanilla model (**c**). Switching thresholds are indicated by the dashed lines (green: hebb → BP, black: BP → hebb). **g** Spectrum of the recurrent weight matrix at initialization same as vanilla model (**d**). **h** Example inputs (green, blue, orange, and gray lines) for two selected tasks, “DelayGo” and “MultiDelayDM”. Shaded areas represent the time period where the loss function is computed. **i** Recurrent activity during the example trials for both model types for each task. **j** Target outputs (black lines) and model responses (green lines) for the example trials for both model types for each task.

Most experimentally observed synaptic plasticity rules are local, relying on the activity of pre-and post-synaptic neurons^14–19^. Local rules are biologically plausible and computationally efficient, and they can support memory and associative learning^14,15,20–25^. Complex behavior, however, also requires credit assignment: error or value information must influence the synapses that contributed to a behavioral outcome^26–31^. Proposed mechanisms for credit assignment in the brain use non-local information to update synapses while preserving biological plausibility, including feedback alignment^32^, dendritic predictive coding^33,34^, and three-factor learning rules^29,35,36^. Such alternatives to the strong, but biologically-implausible, credit assignment that backpropagation offers often fail to scale to more complex tasks^37^. Here, we propose that only a subset of synapses need to access non-local information for successful learning, thus removing a substantial part of the credit assignment computational burden without hurting task learning.

Established computational models addressing synaptic states do not capture the computational role of synaptic heterogeneity. Instead, they typically develop neurally-inspired algorithms that improve learning or memory. For example, such previous models have limited dynamics and learning capacity, generally lacking any type of credit assignment and focusing on distinct synaptic update timescales only^38^, improved learning mechanisms by loading individual synapses with several learning rules^39,40^, or hand-designed specific fixed modules of the network to solve different aspects of the task at hand^41^. Our work is built around connectomic observations that individual synapses may have distinct plasticity states that depend on prior experience of the organism, thus being able to switch between states as needed to maintain a balance between flexible learning and efficient energy consumption.

Our assumptions are also motivated by theory. Individual gradient-based synaptic learning rules typically produce unimodal weight distributions^42,43^, whereas cortical excitatory synapses can show bimodal structure^13^. A bimodal synaptic distribution could therefore reflect more than one learning process operating within the same recurrent circuit. The unresolved question is whether such heterogeneity merely reflects biological detail or whether it changes what and how a circuit can learn, how a circuit computes, and how it affects the circuit’s structure.

We tested this question with a recurrent neural network in which synapses switch between learning rules based on their strength. Weight magnitude is an operational proxy for synaptic state in this model, not a one-to-one mapping onto spine apparatus or any single organelle, and the specific switching thresholds are hyperparameters rather than fitted biological boundaries. Synapses in the low-weight state followed a local Hebbian-like rule, whereas synapses in the high-weight state received an idealized non-local credit-assignment signal implemented with backpropagation through time. This model treats backpropagation through time as a computational proxy for strong credit assignment, not as a proposed biological mechanism. It asks whether state-dependent synaptic switching can reduce the number of recurrent weights that receive BP-based updates while preserving the computational benefits of credit assignment. We found that plasticity-switching RNNs learned cognitive tasks faster than fully backpropagation-trained RNNs, and that switching produced recurrent structural signatures that can be tested in connectomic data. These results provide a set of falsifiable hypotheses for neural circuit structure and suggest a computational role for synaptic structural states: they may control how local plasticity and credit assignment are distributed across a recurrent circuit. Our work, directly motivated by the recently quantified synaptic heterogeneity, provides new insights into how the brain learns using variable components and how these components alter the circuit structure.

## 2 Methods

### 2.1 Network Architecture

We used recurrent neural networks (RNNs) because they provide a simple model of temporal circuit dynamics. The models combine two classes of synaptic updates: local Hebbian-like plasticity and an idealized non-local credit-assignment signal implemented with backpropagation through time (BPTT; for brevity, BP; Fig. 1b,e). We simulated all networks in discrete time (Supplementary Methods). All parameters in the control models (vanilla) were trained using BP. Recurrent weights were initialized from a uniform <u>dist</u>ribution scaled to set the spectral radius at initialization to 1 (Fig. 1c,d), using a scale of 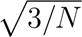 where *N* is the number of hidden units. This initialization is known to provide a good starting point for artificial networks^44–48^. Unless otherwise specified, vanilla RNNs had *N* = 256 hidden units and used the tanh transfer function for the hidden layer and a linear transfer function for the output layer.

Our plasticity-switching RNN (psRNN) shares the same computation and connectivity graph as the vanilla model, but the recurrent weights differ in how they learn (Fig. 1e). We divided recurrent weights into two subsets based on their strength and applied a different learning rule to each subset. A binary dynamic mask **M** switches synaptic learning states between local Hebbian-like plasticity and BP-based credit assignment as synapse weights cross state thresholds.

In line with simple synapses generally having smaller spine heads than spine-apparatus synapses^13^, synapses with small absolute weights use Hebbian-like plasticity, whereas synapses with large absolute weights receive BP updates. This threshold rule is a modeling abstraction. It captures the hypothesis that synaptic state can gate access to different learning signals, but it does not specify the molecular pathway by which a biological synapse would switch state. The threshold values are model hyperparameters and were not fitted to measured synapse-size distributions. Recurrent weights were initialized identically to the vanilla model (Fig. 1f), using the same random seeds for matched comparisons, w<u>ith s</u>witching thresholds set at *θ_BP→Hebb_* = 0.5 and *θ_Hebb→BP_* = 0.8 in units of the network scale 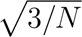 (Fig. 1f, dashed lines). These thresholds made psRNNs start with about 50% of recurrent weights in each subset. The two thresholds differ to account for overlap between synaptic states and to prevent unstable rapid switching. All reported results used the same thresholds unless otherwise noted, although we explored a range of switching thresholds (Supplementary Methods). The spectrum of the recurrent weight matrix at initialization was identical for both model types (Fig. 1g). Unless otherwise specified, psRNNs had *N* = 256 hidden units and used the same transfer functions as the vanilla model.

### 2.2 Tasks

We evaluated both model types on cognitive tasks from the set developed by Yang et al.^49^ for studying task representations in recurrent neural networks. These tasks are widely used in computational neuroscience^50–54^ and require temporal integration, working memory, and context-dependent responses. They therefore provide a useful setting for asking how local Hebbian-like plasticity interacts with non-local credit assignment. We focus on two representative tasks, delayed response (DelayGo)^55^ and multi-sensory delayed decision making (MultiDelayDM)^56^ (Fig. 1h,j; see also Fig. S1a,d for alternative input-target visualizations consistent with prior work). Results for additional non-human-primate-inspired cognitive tasks are reported in Tab. 1, Tab. 2, Tab. S1, and Tab. S2.

**Table 1:**
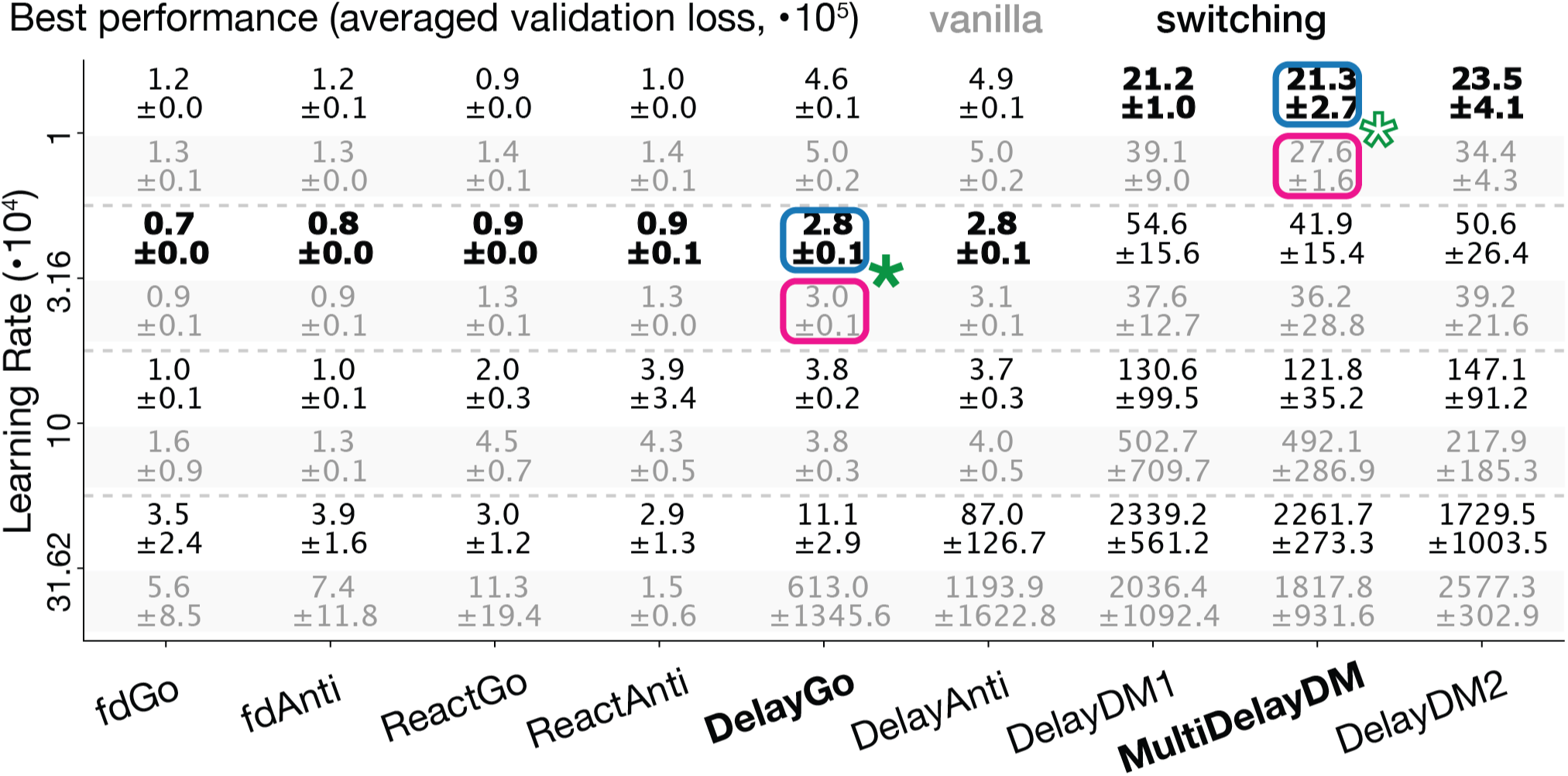
Best performance across tasks and BP learning rates. Mean ± standard deviation of averaged validation loss across 5 random initializations. For robustness, validation loss was averaged across last 10 epochs of training across all validation trials. Vanilla models results are shown in gray and switching models results are shown in black. For each task, the best performance across all BP learning rates is shown in bold. All bold values are from switching models. Green stars link to the corresponding experiments in Fig. 2, highlighted by the pink (vanilla) and blue (switching) boxes. Tasks are ordered by increasing difficulty, i.e. best performance across our trained models, from left to right.

| Best performance (averaged validation loss, $\cdot 10^5$) | | | | | | vanilla | switching | | | |
| --- | --- | --- | --- | --- | --- | --- | --- | --- | --- | --- |
| Learning Rate ($\cdot 10^4$) | 1 | 1.2<br>$\pm 0.0$ | 1.2<br>$\pm 0.1$ | 0.9<br>$\pm 0.0$ | 1.0<br>$\pm 0.0$ | 4.6<br>$\pm 0.1$ | 4.9<br>$\pm 0.1$ | **21.2**<br>**$\pm 1.0$** | **21.3**<br>**$\pm 2.7$** | **23.5**<br>**$\pm 4.1$** |
| | | 1.3<br>$\pm 0.1$ | 1.3<br>$\pm 0.0$ | 1.4<br>$\pm 0.1$ | 1.4<br>$\pm 0.1$ | 5.0<br>$\pm 0.2$ | 5.0<br>$\pm 0.2$ | 39.1<br>$\pm 9.0$ | 27.6<br>$\pm 1.6$ | 34.4<br>$\pm 4.3$ |
| | 3.16 | **0.7**<br>**$\pm 0.0$** | **0.8**<br>**$\pm 0.0$** | **0.9**<br>**$\pm 0.0$** | **0.9**<br>**$\pm 0.1$** | **2.8**<br>**$\pm 0.1$** | **2.8**<br>**$\pm 0.1$** | 54.6<br>$\pm 15.6$ | 41.9<br>$\pm 15.4$ | 50.6<br>$\pm 26.4$ |
| | | 0.9<br>$\pm 0.1$ | 0.9<br>$\pm 0.1$ | 1.3<br>$\pm 0.1$ | 1.3<br>$\pm 0.0$ | 3.0<br>$\pm 0.1$ | 3.1<br>$\pm 0.1$ | 37.6<br>$\pm 12.7$ | 36.2<br>$\pm 28.8$ | 39.2<br>$\pm 21.6$ |
| | 10 | 1.0<br>$\pm 0.1$ | 1.0<br>$\pm 0.1$ | 2.0<br>$\pm 0.3$ | 3.9<br>$\pm 3.4$ | 3.8<br>$\pm 0.2$ | 3.7<br>$\pm 0.3$ | 130.6<br>$\pm 99.5$ | 121.8<br>$\pm 35.2$ | 147.1<br>$\pm 91.2$ |
| | | 1.6<br>$\pm 0.9$ | 1.3<br>$\pm 0.1$ | 4.5<br>$\pm 0.7$ | 4.3<br>$\pm 0.5$ | 3.8<br>$\pm 0.3$ | 4.0<br>$\pm 0.5$ | 502.7<br>$\pm 709.7$ | 492.1<br>$\pm 286.9$ | 217.9<br>$\pm 185.3$ |
| | 31.62 | 3.5<br>$\pm 2.4$ | 3.9<br>$\pm 1.6$ | 3.0<br>$\pm 1.2$ | 2.9<br>$\pm 1.3$ | 11.1<br>$\pm 2.9$ | 87.0<br>$\pm 126.7$ | 2339.2<br>$\pm 561.2$ | 2261.7<br>$\pm 273.3$ | 1729.5<br>$\pm 1003.5$ |
| | | 5.6<br>$\pm 8.5$ | 7.4<br>$\pm 11.8$ | 11.3<br>$\pm 19.4$ | 1.5<br>$\pm 0.6$ | 613.0<br>$\pm 1345.6$ | 1193.9<br>$\pm 1622.8$ | 2036.4<br>$\pm 1092.4$ | 1817.8<br>$\pm 931.6$ | 2577.3<br>$\pm 302.9$ |
|  |  | fdGo | fdAnti | ReactGo | ReactAnti | DelayGo | DelayAnti | DelayDM1 | MultiDelayDM | DelayDM2 |

**Table 2:**
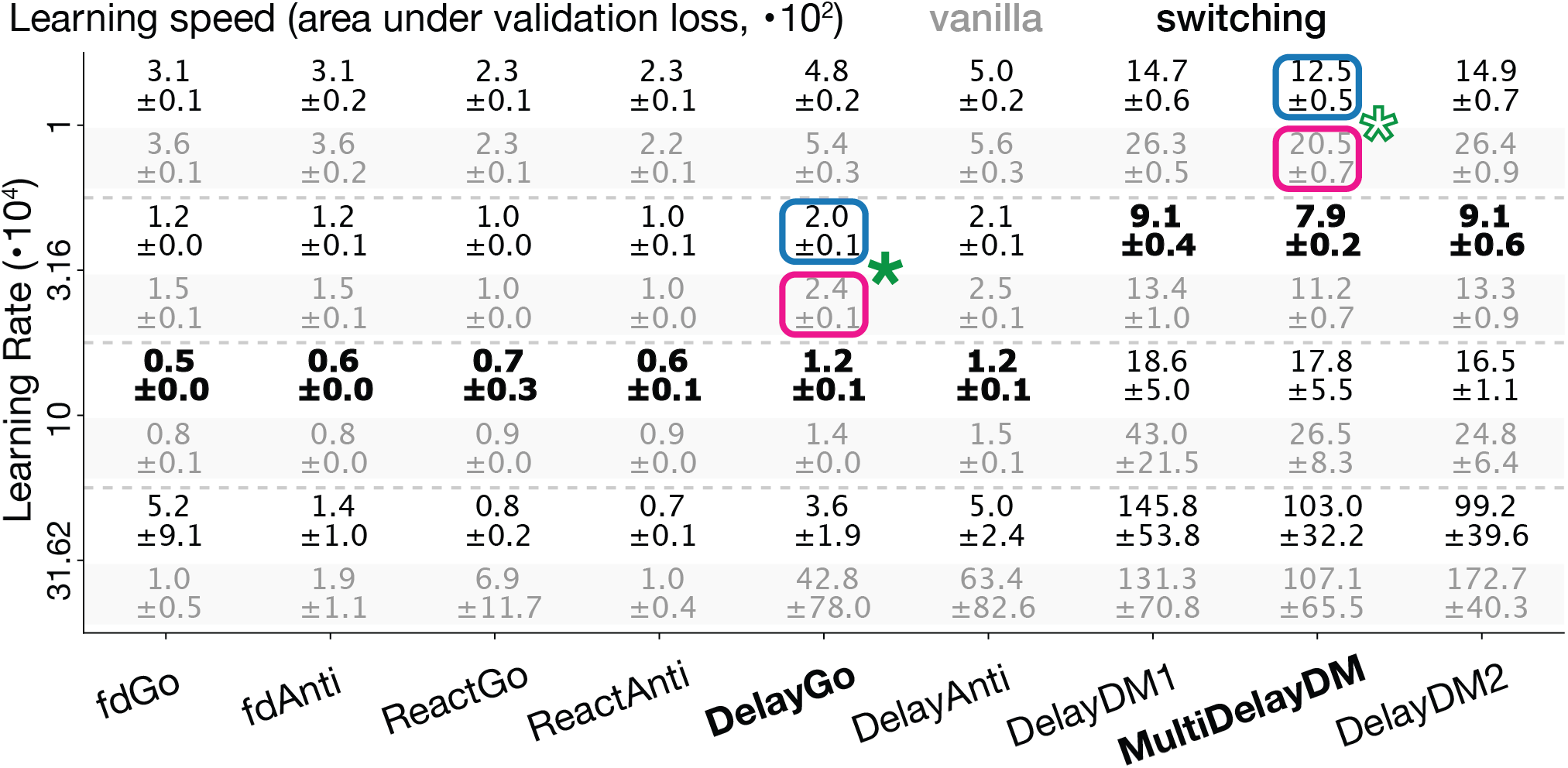
Learning speed across tasks and BP learning rates. Mean ± standard deviation of learning speed across 5 random initializations. Learning speed is defined as the area under the curve of validation loss throughout training. All other details are the same as in Tab. 1. Tasks maintain same order as in Tab. 1.

The DelayGo task (moderate difficulty) requires the network to maintain a spatial location in working memory during a delay period and produce a response at the remembered location after a go cue^55^. The MultiDelayDM task (high difficulty) extends this challenge by requiring comparison between multiple stimuli presented sequentially with varying delay periods, placing additional demands on working memory (extension of the task from Raposo et al.^56^). Both tasks feature distinct temporal phases (stimulus presentation, delay, and response) that provide natural opportunities for Hebbian plasticity to operate on correlated activity patterns while simultaneously requiring precise credit assignment for successful performance.

### 2.3 Training

All parameters shown in pink in Fig. 1b and Fig. 1e receive BP updates. Parameters shown in blue in Fig. 1e are updated using a local Hebbian-like rule with decay *δ* and parameter-specific BP-trainable learning rate **Λ**. The update rule is given by **W***_Hebb_* ← *δ* · **W***_Hebb_* + **Λ** ⊙ Δ**W***_Hebb_*, where ⊙ denotes element-wise multiplication. The Hebbian update Δ**W***_Hebb_* is computed as 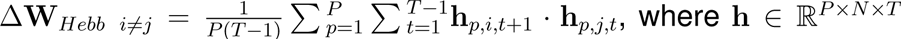 (number of trials included in Hebbian plasticity *P*, hidden units *N*, time steps *T*) is the hidden activity over the last *P* trials. To avoid unstable self-amplification, we set the diagonal of the Hebbian update to zero, Δ**W***_Hebb i,i_* = 0. To allow BP to account for Hebbian updates, we set the Hebbian period *P* smaller than the batch size *B* and took multiple Hebbian steps during one BP batch. Unless otherwise specified, psRNNs took 2 Hebbian steps per BP batch, with *P* = 128 = *B/*2 trials per period.

We explored several regimes for the Hebbian learning rate. In the default configuration, the Hebbian learning rate, **Λ**, was BP-trained independently for each Hebbian synapse. This choice gives the vanilla and switching models the same number of BP-trained parameters, making the comparison more conservative. BP-training individual synaptic learning rates is a computational simplification. In biological circuits, heterogeneous plasticity parameters could arise from evolutionary, developmental, or active selective control processes rather than gradient-based optimization^57–61^. We therefore treated psRNN variants with fixed Hebbian learning rates and many fewer trained parameters as an important control on the role of meta-optimized plasticity; these variants also perform well (Fig. S2b,c). We tested several choices for the decay parameter *δ*, including heterogeneous and BP-trained variants, but these did not substantially affect performance. We therefore set *δ* = 0.9999 throughout all experiments. **Λ** was initialized at 0 for all Hebbian synapses to remove any initial Hebbian bias.

We verified that both RNNs and psRNNs produce similar dynamics after training (Fig. 1i). Internal dynamics are projected to the output space via the output weights and biases, which are BP-trained parameters in both model types. Both model types eventually solve the task and produce behavior that follows the target (Fig. 1j). For alternative visualizations of the hidden activity and responses, consistent with prior work using these tasks, see Fig. S1b,c.

## 3 Results

### 3.1 State-dependent plasticity improves learning despite reduced synaptic credit assignment

We first asked whether a recurrent network can learn well when many recurrent weights do not receive BP-based updates. We trained vanilla RNNs and psRNNs on DelayGo and MultiDelayDM using the best BP learning rate for each model type and task, selected from a sweep across a range of values (Fig. S2a). By design, many recurrent weights in psRNNs use only the Hebbian-like rule until, if ever, they cross into the BP subset. Nevertheless, psRNNs reached lower training and validation loss than vanilla RNNs across tasks (Fig. 2a,b; Tab. 1). They also learned faster, as measured by the area under the validation loss curve (Fig. 2a,b; Tab. 2). These results generally held across network sizes (Tab. S1, Tab. S2) and transfer functions (Tab. S3). Behavioral performance, measured as validation accuracy, was also higher for psRNNs even though accuracy was not directly optimized (Fig. 2c; see Supplementary Methods). These comparisons represent model-level replication across random initializations and hyperparameter sweeps, and we report uncertainty across seeds in figures and tables.

**Figure 2:**
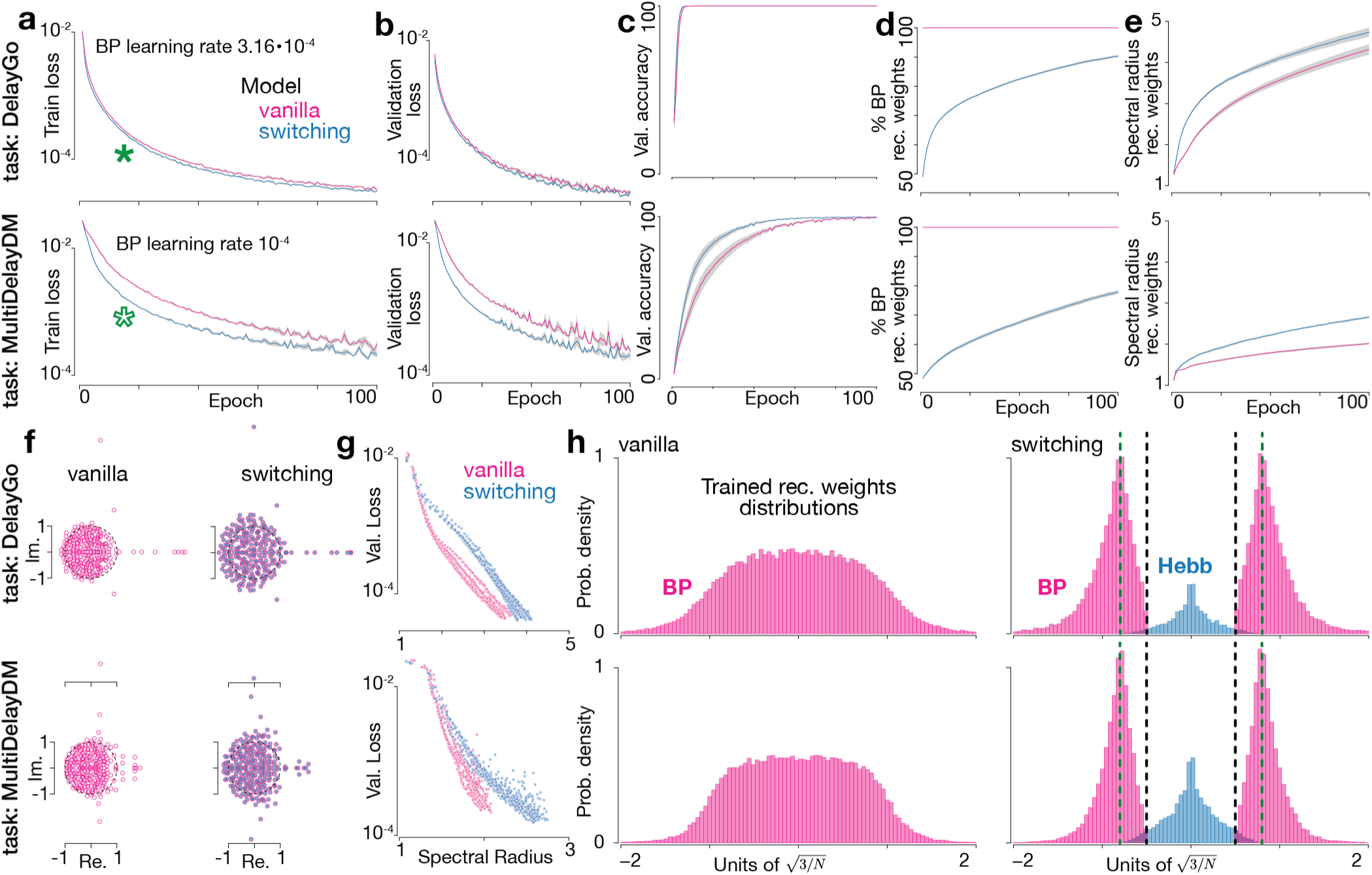
Switching plasticity improves learning while reducing the recurrent synapses receiving credit assignment. **a** Training loss as a function of training epochs for both model types and tasks. Shaded areas represent standard error of the mean (s.e.m.) across 5 random initializations. Green stars link to the corresponding experiments in Tab. 1 and Tab. 2. BP learning rate for shown experiments was chosen based on a parameter sweep across different values (Fig. S2a) to optimize the performance of each model type individually, while maintaining stability throughout training. **b** Validation loss, **c** validation accuracy, **d** percent BP recurrent weights, and **e** spectral radius or recurrent weights throughout training, otherwise same as **a**. Accuracy computation is described in the Supplementary Methods section. **f** Example spectrum of the recurrent weight matrix after training for both model types and tasks. Dashed lines indicate the unit circle. **g** Validation loss as a function of the spectral radius for all models for each task throughout training. Individual points represent a single random initialization at a specific training epoch. **h** Example trained recurrent weights distribution for the vanilla (left) and switching (right) model for each task. Long tails of distributions are truncated at 2 in units of network scale for visualization purposes. Switching thresholds (Fig 1f) are shown on the switching model weights distribution.

The performance gain was not caused by giving psRNNs more access to credit assignment. psRNNs used fewer BP-updated recurrent weights than vanilla RNNs throughout training (Fig. 2d), especially during the early learning period when the performance gap emerged. In the default model, BP also optimizes the Hebbian learning-rate matrix **Λ**, so this result should be interpreted as reduced access to credit assignment for recurrent weights rather than complete removal of gradient information from every Hebbian synapse. When BP optimized the Hebbian learning rates, psRNNs gradually moved more synapses into the BP subset. However, synapses switched in both directions rather than simply accumulating in the BP subset (Fig. S2d). The model therefore maintained an ongoing interaction between local Hebbian-like plasticity and non-local credit assignment.

psRNNs also changed recurrent dynamics during learning. The spectral radius of the recurrent weight matrix increased faster in psRNNs than in vanilla RNNs (Fig. 2e), potentially allowing longer timescales through larger real eigenvalues and faster input responses through larger complex eigenvalues. This difference is visible in the trained spectra (Fig. 2f), where psRNNs show more eigenvalues with large real and imaginary components. Across models and training epochs, larger spectral radius was associated with lower validation loss (Fig. 2g). Switching also produced a multimodal recurrent weight distribution after training (Fig. 2h), qualitatively matching the bimodal synaptic weight distributions observed in cortical connectomic data^13^ and the theoretical expectation that a single learning rule tends to produce unimodal synaptic weights^42^.

Several simpler psRNN variants also performed well, reaching performance comparable to or slightly better than vanilla RNNs (Fig. S2b and Supplementary Methods). This control is important because the default model uses a BP-trained Hebbian learning-rate matrix. The best-performing variant used heterogeneous BP-trained Hebbian learning rates (Λ is a BP-trained matrix), but simpler variants fixed the Hebbian learning rates to heterogeneous (Λ is a fixed matrix) or homogeneous (Λ is a fixed scalar) values. Thus, BP-based optimization of Λ improves the model but is not necessary for state-dependent switching to retain good performance. We also removed both Hebbian plasticity and switching to create a semi-reservoir baseline in which the smallest ∼ 50% of recurrent synapses at initialization were frozen and the rest received BP updates. This baseline connects to classical reservoir computing^62–64^ and performed close to vanilla RNNs, showing that fully training every recurrent weight is not necessary for these tasks. Nevertheless, the psRNN variants surpassed vanilla and semi-reservoir baselines in performance (Fig. S2b), credit-assignment efficiency (Fig. S2c), or both.

Variants with active Hebbian-like learning showed that the non-BP subset contributes task-relevant updates. Depending on how Λ was constrained, Hebbian-like plasticity either improved performance while moving more synapses into the BP subset (fixed or optimized Λ matrix) or preserved performance while shrinking the BP subset (fixed Λ scalar). When BP could not optimize Hebbian learning rates, the percentage of BP recurrent weights saturated at lower values (Fig. S2c, compare orange and green curves to blue curve), with strong performance even when only about 35% of recurrent weights belonged to the BP subset.

### 3.2 Several plasticity interactions are responsible for improved learning

The performance advantage of psRNNs arose from the interaction between local plasticity, credit assignment, and switching. First, Hebbian updates occurred multiple times within each BP batch, allowing BP to evaluate multiple nearby parameter configurations per gradient step (Fig. 3a). Second, Hebbian-like plasticity reshaped recurrent connectivity from the activity generated by each task, creating a dynamic and task-specific initialization for later BP updates (Fig. 3b). Third, switching prevented Hebbian-like updates from pushing synapses into unfavorable regions of parameter space and moved those synapses into the credit-assignment subset while they remained useful (Fig. 3c,d and S3).

**Figure 3:**
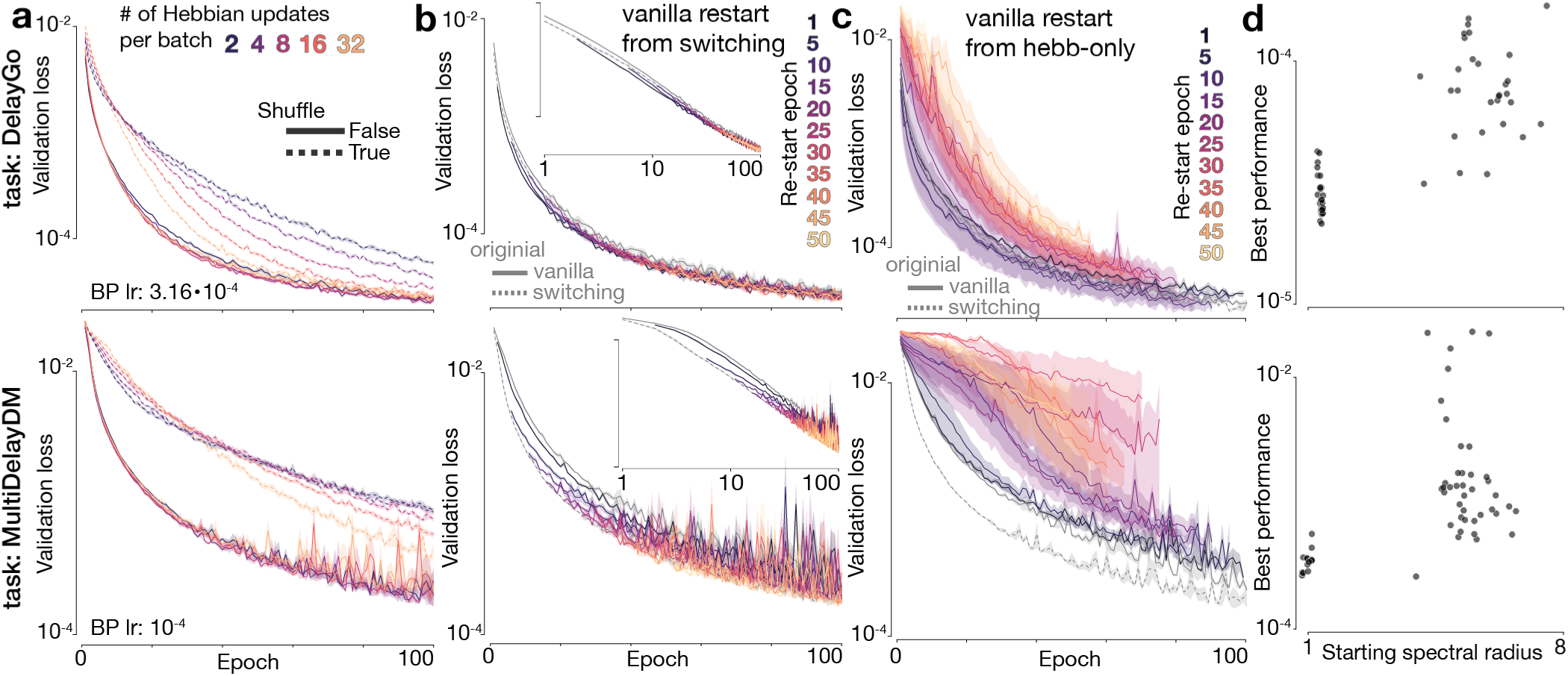
Local plasticity improves learning only when coordinated with credit assignment and switching. **a** Validation loss throughout training for switching models for each task as a function of the number of Hebbian steps during one BP batch (solid lines, color-coded). Dashed lines represent controls where the same number of weights update steps are taken during one BP batch, but the Hebbian updates are shuffled onto the Hebbian synapses. Shaded areas represent s.e.m. across 10 random initializations. **b** Validation loss throughout training for vanilla models (completely BP-trained) for each task, initialized with the same weights from pre-trained switching models, color-coded by the number of pre-trained epochs of the switching model used for initialization (solid colored lines). Gray lines represent the original vanilla (solid) and switching (dashed) models for reference. Plotted validation loss for the vanilla models starts at the corresponding number of pre-trained epochs. Shaded areas represent s.e.m. across 5 random initializations. Insets show the same data in log-log plot to better visualize the filled gap between the original models. **c** Similar to **b**, but initialization is done based on Hebb-only pre-trained models. Since no credit assignment is used during pre-training, plotted validation loss for the vanilla models starts at epoch 0 for all numbers of Hebb-only pre-trained epochs. **d** Best performance of the vanilla models in **c** as a function of the spectral radius of the pre-trained Hebb-only models used for initialization. Best performance computed as in Tab. 1. Each point represents a single random initialization.

To test the contribution of multiple parameter-space samples, we varied the number of Hebbian updates taken within each BP batch. Throughout the main results, psRNNs used 2 Hebbian updates per BP batch, the minimum needed for BP to account for the Hebbian updates with-out substantially increasing training cost. Increasing the number of Hebbian updates slightly improved performance up to a task-dependent point, after which excessive updates started to destabilize training (Fig. 3a). This parameter-space sampling benefit was more prominent in our control experiments: when we shuffled Hebbian updates across Hebbian synapses, removing the correct temporal correlations, performance still improved with the number of updates (Fig. 3a, dashed lines). The shuffled control nevertheless performed substantially worse than the original model, showing that the task-relevant structure of the Hebbian updates is crucial.

The Hebbian-like component also generated a dynamic, task-relevant initialization for the BP component. To isolate this effect, we restarted vanilla RNNs from psRNN checkpoints collected throughout training while matching the initial random seed (Fig. 3b). Vanilla RNNs initialized from psRNN checkpoints closed the performance gap between randomly initialized vanilla RNNs and psRNNs, and after enough pre-training they slightly surpassed psRNNs. The required amount of pre-training varied by task, but even 1 epoch of psRNN pre-training significantly improved vanilla RNN performance (Fig. 3b). This result suggests that Hebbian-like updates are most useful early in learning, when they move the recurrent network into a region of weight space that BP can then exploit more efficiently.

Hebbian-like learning alone did not explain the benefit. To test the role of the dynamic switching interaction on improved learning in our models, we trained “Hebb-only” psRNNs by setting the switching thresholds to ∞, preventing recurrent synapses from receiving BP updates. Vanilla RNNs restarted from these Hebb-only checkpoints generally performed worse than networks started from random initialization (Fig. 3c). DelayGo showed slight improvement with minimal Hebb-only pre-training in some cases, but this effect did not generalize across tasks and was highly sensitive to pre-training duration. These results show that the benefit of psRNNs does not come from local plasticity in isolation. It depends on switching, which allows local plasticity to shape useful weights while preventing unconstrained Hebbian growth from dominating the recurrent matrix.

The spectral radius provides one view of why switching is needed. Proper initialization scaling is known to be critical for training recurrent and feedforward networks^44–48^. In vanilla RNNs restarted from Hebb-only checkpoints, the starting spectral radius strongly predicted the final performance of the restarted network (Fig. 3d). Individual Hebb-only models often showed abrupt increases in spectral radius during training (Fig. S3a), and these increases coincided with poor learning after restart (Fig. S3b). Thus, Hebbian-like learning can create useful task-dependent structure, but switching is needed to keep that structure within a range where credit assignment remains effective.

### 3.3 Switching models predict distinctive recurrent circuit structure

If synaptic state routes plasticity, it should change not only learning speed but also the structure of the trained recurrent circuit. We therefore compared the recurrent weight matrices of psRNNs and vanilla RNNs after and during training. We present results for both tasks with *N* = 256 hidden units, and the findings held across all network sizes tested (Fig. S4). These structural differences provide a set of principled and falsifiable hypotheses that could be tested in connectomic graphs, based on our switching-plasticity premise.

Using a metric based on the symmetric and skew-symmetric components of the recurrent weight matrix^65^ (see Supplementary Methods), we found that psRNNs developed more anti-symmetric recurrent weights than vanilla RNNs (Fig. 4a). The departure from 0 was small, but it was consistent across tasks, network sizes, and random seeds (Figs. 4a & S4a). This anti-symmetric shift is consistent with the faster growth of complex eigenvalues in psRNNs and with prior work linking anti-symmetric connectivity to learning and stable memories under some homeostatic mechanisms^66,67^. The anti-symmetry emerged from the interaction between learning rules. Although the Hebbian update is nearly symmetric, both synapse subsets became more anti-symmetric during training, and the BP subset became more anti-symmetric than the Hebbian subset (Fig. 4b). This result suggests that the Hebbian subset changes how credit assignment shapes the synapses that receive BP updates.

**Figure 4:**
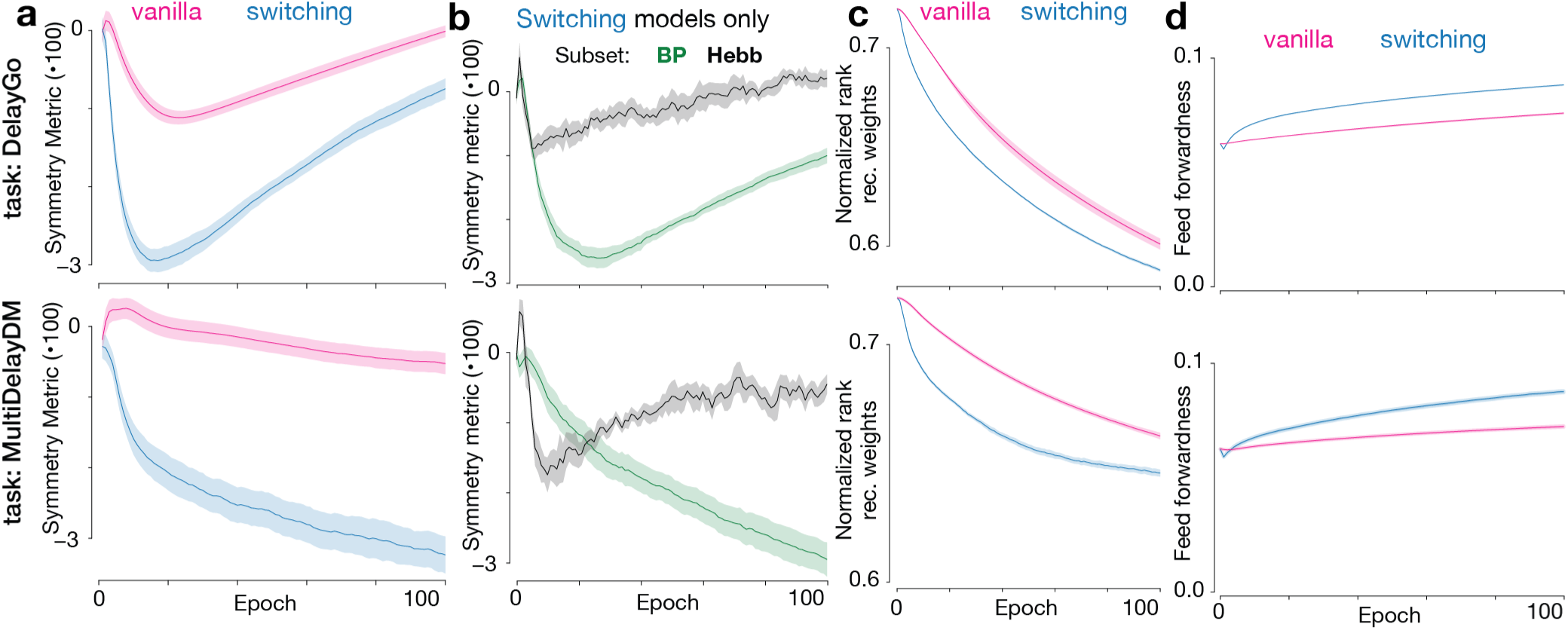
Switching plasticity produces structural signatures. **a** Symmetry metric of the recurrent weights matrix, **b** symmetry metric of the Hebb and BP subsets of the recurrent weights matrix for switching models, **c** normalized rank of the recurrent weights matrix, and **d** amount of feed forward structure in the recurrent weights matrix, all plotted as a function of training epochs for both model types and tasks. Symmetry metric, normalized rank, and feed forwardness computations are described in the Supplementary Methods section. Shaded areas represent s.e.m. across 5 random initializations.

psRNNs also developed lower effective rank. Low-rank structure is important in both neuroscience and machine learning because low-dimensional tasks are often solved by large neural populations^68–79^. Prior work has typically studied low-rank networks by imposing explicit architectural constraints^69,71–75,77,78^. We instead measured whether lower-rank structure emerges from plasticity switching. Using the participation ratio of the recurrent singular values (see Supplementary Methods), we found that psRNNs developed lower normalized rank than vanilla RNNs (Fig. 4c). The trained matrices were not low-rank in the strict sense, but switching moved the network toward lower effective dimensionality without imposing a low-rank constraint.

Finally, psRNNs developed stronger feedforward structure within the recurrent matrix. Recurrent circuits can rely on both feedback loops and feedforward pathways, and the balance between these components shapes circuit computation^80–90^. We quantified feedforward structure using Henrici’s departure from normality^91^, normalized by network size (see Supplementary Methods). More non-normal recurrent matrices contain stronger feedforward components in their Schur decomposition. psRNNs developed larger normalized departure from normality than vanilla RNNs (Fig. 4d), suggesting that these circuits may rely more on feedforward pathways within their recurrent structure.

These predictions can be tested in connectomic graphs by asking whether synapse classes associated with different structural states also differ in symmetry, effective rank, or feedforward organization. A direct test would separate recurrent excitatory connections by spine-apparatus annotation or by synaptic size proxy in lack of well-annotated synapses, compute structural metrics in the circuit as a whole and within each group, and compare the observed values against degree-, weight-, and spatially-matched null graphs. Our model predicts that neural circuits involving multiple state-defined synaptic groups should be different from control circuits in their main computational pathway type and effective dimensionality. Conversely, the structural hypothesis would be weakened if well-annotated connectomic datasets showed no state-dependent differences after matching for weight, degree, spatial proximity, and cell-type composition.

## 4 Discussion

We introduced psRNNs to test whether synaptic state can regulate how local plasticity and credit assignment are distributed across a recurrent circuit. In these models, weak synapses follow a Hebbian-like rule, whereas strong synapses receive an idealized non-local credit-assignment signal implemented with BP. This simple state-dependent rule improved learning across cognitive tasks even though fewer recurrent weights received BP-based updates. The benefit depended on the interaction between local plasticity, credit assignment, and switching: Hebbian-like updates created task-relevant recurrent structure, BP exploited that structure, and switching prevented unconstrained Hebbian growth from destabilizing the network.

This framework offers a computational interpretation of synaptic structural heterogeneity. Experimental studies often observe local plasticity rules, but these rules may represent only the first-order terms of richer synaptic update rules that include non-local components^38,92,93^. We considered a complementary possibility: different synapses may use different learning rules depending on their state. Spine-apparatus synapses are a natural motivation for this idea because they differ from smaller synapses in ultrastructure and calcium handling^1–4^. Our results suggest that this kind of state dependence could do more than reflect synaptic history. It could help a recurrent circuit decide where costly or specialized credit-assignment signals are most useful.

The model is intentionally idealized. BP provides a strong credit-assignment signal, but it is not a biologically realistic mechanism in its standard form. We therefore interpret BP here as an upper-bound proxy for non-local information that could be carried by more plausible mechanisms, such as three-factor learning rules, dendritic credit-assignment signals, or eligibility-trace-based approximations^29,33–35^. The Hebbian-like updates also operate on longer timescales than many biological plasticity processes and omit short-term dynamics typically used in other computational models^39,40,65^. The switching thresholds reduce complex molecular and structural synaptic states to a scalar weight rule and were chosen as computational hyperparameters, not inferred from biological synapse classes. Finally, the default psRNN optimizes the Hebbian learning rate at each synapse using BP, which is a computational convenience rather than a proposed biological process. Simpler variants with fixed Hebbian learning rates still performed well (Fig. S2b,c), suggesting that meta-learned plasticity improves learning speed and final performance but is not required for the switching principle to work. The current experiments therefore support the principle that state-dependent rule assignment can be useful, but do not identify the biological mechanism that implements credit assignment.

The strongest experimental implication of psRNNs is structural. Switching plasticity altered recurrent symmetry, effective rank, and feedforward structure, producing signatures that can be measured in connectomic graphs. Symmetry and anti-symmetry are related to attractor structure, memory, and stability^67,94,95^; effective rank reflects how many dimensions of the population are used for computation^68,69,71^; and feedforward structure captures directed pathways embedded within recurrent connectivity^80,82,83,86–88^. These measurements can be applied to dense model matrices and sparse connectomic graphs. The effects in our models are modest, especially for the symmetry metric, so detecting them in biological data will require sufficiently large and well-annotated synapse classes. Nevertheless, the predictions are falsifiable: synapses in different structural states should differ not only in size or organelle content, but also in their contribution to recurrent circuit organization.

psRNNs also connect to machine-learning ideas, but the analogy is secondary to the circuit hypothesis. The switching mechanism resembles an online form of sparse training or network pruning, because it dynamically identifies which recurrent weights receive BP-based updates. It also relates to reservoir computing, where a fixed recurrent layer is paired with a trained read-out^62–64^. In psRNNs, however, the recurrent layer is neither fully fixed nor fully trained by BP. Local plasticity continuously changes part of the recurrent network, while switching determines which synapses join the credit-assignment subset. This hybrid organization may explain why psRNNs can outperform fully BP-trained RNNs during learning rather than merely match them with fewer trainable parameters.

Several extensions would test the biological scope of the switching principle. The most direct computational test is to replace BP with biologically plausible approximations, such as three-factor learning rules, and ask whether the benefit persists when credit assignment is noisy, delayed, or incomplete^29,35^. That experiment is beyond the present study, so the current results establish the utility of selective access to a strong teaching signal rather than the sufficiency of any particular biological approximation. Adding Dale’s law and cell-type identity, for example with exponentiated gradients^43^, would make the recurrent architecture closer to cortical circuits. Replacing fixed switching thresholds with state variables tied to calcium dynamics, synaptic history, or spine-apparatus state would make the switching mechanism more biological. Testing motor control, navigation, continual learning, or multi-task settings would also show whether switching helps beyond the cognitive tasks studied here.

Our results support a narrower claim: in an idealized recurrent model, local plasticity and a strong teaching signal need not be assigned uniformly across a network. When coordinated by synaptic-state switching, simple local plasticity can improve learning while reducing the number of recurrent weights that receive non-local updates. This proof-of-principle model, broadly focused on the computational role of synaptic heterogeneity rather than algorithms specifically designed for certain tasks, motivates the biological hypothesis that synaptic structural states may route learning signals and yields connectomic predictions for testing whether different synapse classes play different roles in recurrent circuit organization.

## Acknowledgements

We thank the Mihalas group, including Kyle Aitken, Yuan Gao, Lukasz Kusmierz, Zhixin Lu, Dana Mastrovito and Zihan Zhang, and Blake Richards and Roman Pogodin for helpful discussions. DT and SD were supported by the Shanahan Family Foundation Fellowship at the Interface of Data and Neuroscience at the Allen Institute and the University of Washington, supported in part by the Allen Institute. SM was in part supported by NSF (Nos. 2223725 and 2424124) and NIH (Nos. R01EB029813 and RF1DA055669) grants. This research was supported by the Allen Institute, founded by Jody Allen – chair and co-founder of Allen Family Philanthropies, and the late Paul G. Allen – investor, philanthropist, and co-founder of Microsoft. We gratefully acknowledge their vision and generosity, which make this work possible.

## 5 Supplementary material

### 7.0.1 Code availability

All code developed for this work is publicly available at github.com/DenisTurcu/psRNN.

### 7.0.2 LLMs use

All code for the models we used, including control and custom architecture switching models, was designed and written by us. All code for creating the data for all tasks was lightly adapted from Yang et al.^49^ to fit our framework and packages. After obtaining preliminary results, we used “Claude Sonnet 4” and “Claude Sonnet 4.5” to organize the code and to efficiently setting up Python scripts for submitting parallelized Slurm jobs to take full advantage of the compute cluster at Allen Institute. After drafting and revising the first version of the full manuscript, we used “Claude Sonnet 4.6”, “Claude Opus 4.6”, and “GPT-5.5” to point to grammatical inconsistencies, suggest clarity improvements, and improve the flow of the manuscript as it relates to transitions between paragraphs and sections. No additional language models were used after evaluating all suggestions and changes as described above. Finally, we took several individual reviewing rounds before we converged to the last manuscript version.

### 5.1 Supplementary Methods

#### 5.1.1 RNN dynamics

We simulated all rate-based recurrent neural networks in discrete time according to the following equations:

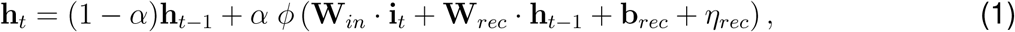

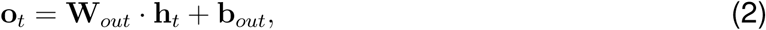

where **h***_t_,* **o***_t_,* **i***_t_* are the hidden state, output, and input at time *t*, *η_rec_* is recurrent noise independent for each neuron (Gaussian with fixed standard deviation), **b***_rec_,* **b***_out_* are the recurrent and output biases, 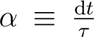 is the dimensionless simulation time step, and *ϕ*(·) is the activation function, specified in the text for each experiment. The weight matrices **W***_in_*, **W***_rec_*, and **W***_out_* have sizes *N_rec_*×*N_in_*, *N_rec_*×*N_rec_*, and *N_out_*×*N_rec_*, respectively. We used standard values for the training input noise (standard deviation 0.01), recurrent noise (standard deviation 0.05), and *α* = 20ms*/*100ms = 0.2, along with all other simulation hyperparameters from p<u>revious st</u>udies^49,53^. We scaled all noise based on the simulation time step *α*, multiplying by 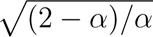, for consistency with the continuous-time limit of the dynamics.

#### 5.1.2 Switching thresholds

Switching thresholds impact both the performance of psRNNs and the final percentage of BP-updated recurrent weights, making them important hyperparameters rather than biological measurements. We swept across a range of values but focus on the best-performing pair for the ma<u>in resu</u>lts: *θ_Hebb→BP_* = 0.8 and *θ_BP→Hebb_* = 0.5 (or (0.5, 0.8) as a pair) in units of network scale 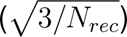. We primarily explored larger thresholds to confirm that psRNNs can solve tasks even when the BP subset is smaller than the Hebbian subset. We ran networks with thresholds {(0.5, 1.0), (0.5, 1.2), (0.7, 1.0), (0.8, 1.1), (0.9, 1.2), (1.0, 1.1)}. In all cases, psRNNs reached 100% validation accuracy within the 100 training epochs and the validation loss started to plateau, despite slow continual decrease. psRNNs with large thresholds solved the task with approximately 35–40% of the recurrent weights in the BP subset by epoch 100, despite starting with 0% or 10% BP-updated recurrent weights at initialization. These models required approximately 5–10 additional epochs to train, as they needed sufficient exposure before receiving meaningful credit assignment through BP.

#### 5.1.3 BP-trained parameters

We trained all BP-governed parameters using BPTT on the mean squared error (MSE) loss between the network output and the target output, weighted by a time-dependent mask (Fig. 1h,j), following Yang et al.^49^. We used the Adam optimizer^96^ for 100 epochs with 500 batches per epoch and *B* = 256 trials per batch. Training data included a small amount of Gaussian noise in each input channel (Supplementary Methods). Validation data was noise-free and contained 8192 trials not used during training. We swept across a range of BP learning rates to optimize the performance and learning speed of each model type individually, while maintaining stability throughout training (Fig. S2a; Tabs. 1, 2, S1, S2, S3). We specify the BP learning rate for all experiments in the corresponding figure panels. All parameters shown in pink in Fig. 1b and Fig. 1e receive BP updates.

#### 5.1.4 Accuracy computation

Although model performance is primarily measured by the training loss, these tasks also offer a cognitive performance metric, based on behavioral accuracy measured across a large number of trials. A trial is labeled correct if (1) the model response remains appropriately silent during fixation, (2) the model fixation output matches the fixation target (high during fixation, silent during response), and (3) the model response is active in the correct response direction according to a population coding vector^49^ and produces a response of sufficient magnitude, all throughout the duration of the trial when the mask is active (Fig. 1h). Margins of error for each criterion were defined based on model responses throughout training and were tighter than in prior work to better capture the performance differences between psRNNs and vanilla RNNs. Specifically, we allowed the model fixation output to vary by a maximum of 0.15 (0.25) during the fixation (response) period, allowed for a maximum angle deviation of 10*^◦^*in the response direction, and required the response magnitude to be within 30% of the target response magnitude.

#### 5.1.5 psRNN variants

The psRNN architecture is flexible and can be implemented in various ways based on very few hyperparameters. We explored several variants of psRNNs to test the importance of different components of the architecture and to demonstrate that the performance advantage of psRNNs is not dependent on specific design choices. For example, we tested variants with fixed Hebbian learning rates. For these variants, we selected values based on the distribution of BP-trained Hebbian learning rates from the best-performing psRNNs (Fig. S2e). Fixed heterogeneous parameters were sampled from the corresponding distribution, and the fixed homogeneous parameter was computed as the standard deviation of the BP-trained Hebbian learning rates divided by 100. Both positive and negative fixed homogeneous learning rates performed well, with moderately better performance for the negative value (results for negative value reported in Fig. S2b,c).

#### 5.1.6 Symmetry metric

We computed the symmetry metric following previous work^65^. The metric is based on the symmetric and skew-symmetric parts of a square matrix **W**:

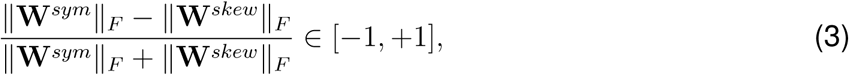

where **W***^sym^* = (**W** + **W***^T^*)*/*2, **W***^skew^* = (**W** − **W***^T^*)*/*2, and ||·||*_F_* is the Frobenius norm. A value of +1 indicates a fully symmetric matrix, −1 indicates a fully anti-symmetric matrix, and 0 indicates balanced symmetric and anti-symmetric components.

#### 5.1.7 Recurrent rank

We used the singular values of the recurrent weight matrix to compute the normalized rank, defined based on the participation ratio as:

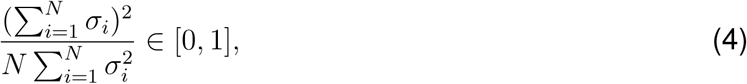

where 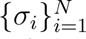 are the *N* singular values and *N* is the number of neurons. A value of 1 indicates a full-rank matrix and a value near 0 indicates a nearly null matrix. Division by *N* normalizes across different network sizes.

#### 5.1.8 Feed forwardness

The Schur decomposition of a square matrix **W** = **Q**(**D** + **T**)**Q***^†^*, where **Q** is unitary, **D** is diagonal containing the eigenvalues, and **T** is strictly upper triangular, and *^†^* represent the conjugate transpose, provides a useful tool for analyzing feed forward structure in recurrent weight matri-ces^97^. All elements of **T** represent feed forward connections and their strengths. The full Schur decomposition, while useful for detailed analysis, is computationally expensive and not required for our purposes. Instead, Henrici’s departure from normality^91^, 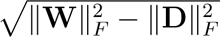, suffices because 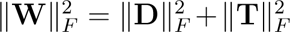 since **Q** is unitary and the Frobenius norm is unitarily invariant. We interpret Henrici’s departure from normality, scaled by the square root of the number of upper triangular elements in the matrix to compare across network sizes, as the amount of feed forward structure of the network:

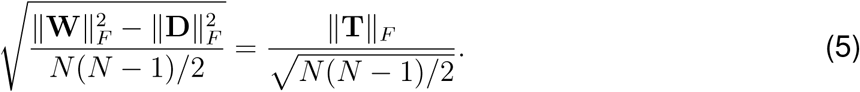

### 5.2 Supplementary figures and tables

**Figure S1:**
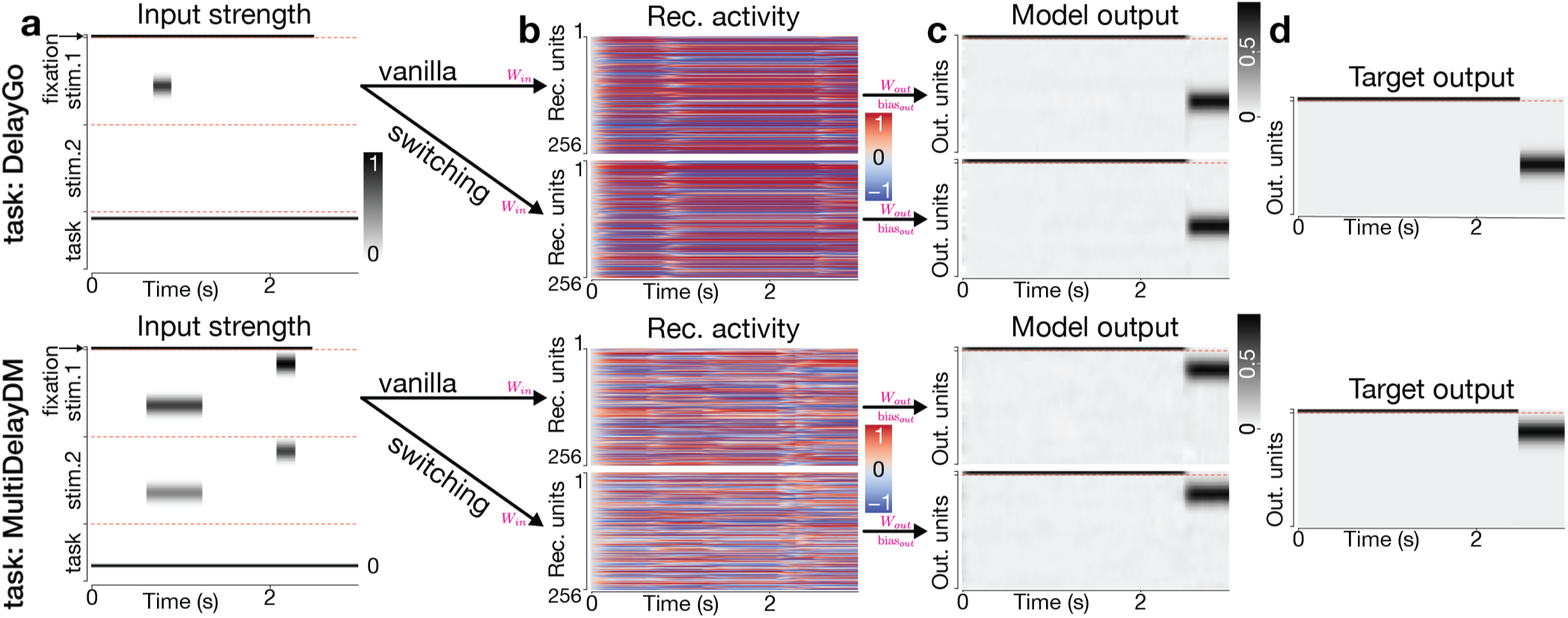
Behavioral dynamics of vanilla and switching models. **a** Example inputs from Fig. 1h, **b** example hidden activity from Fig. 1i, and **c & d** example model responses and targets from Fig. 1j, visualized as heatmaps, in accordance with previous work introducing and using these tasks, for visual reference to these prior studies. The red dashed lines separate the various kinds of input or output dimensions.

**Figure S2:**
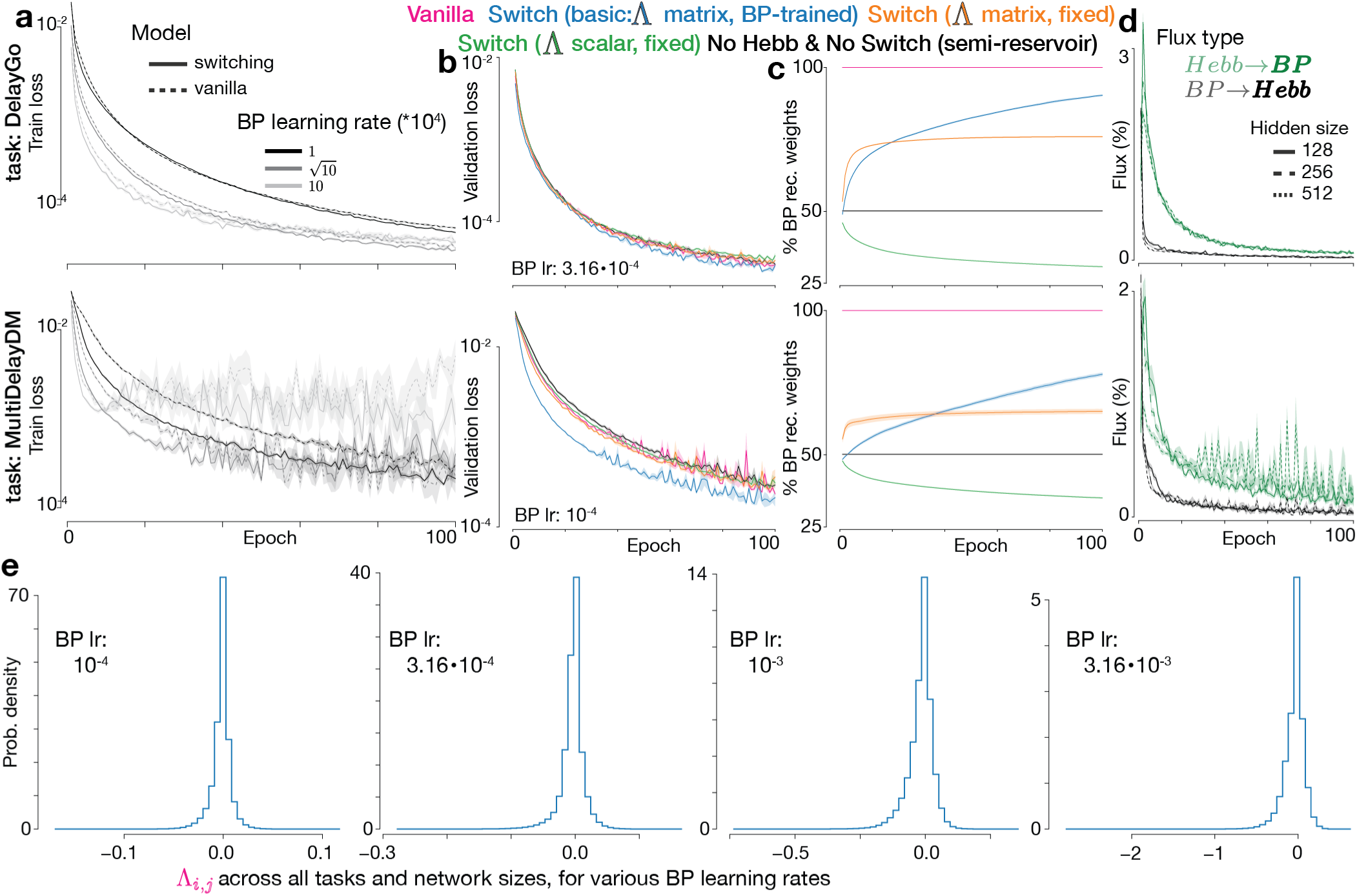
Extended learning dynamics and properties, and switching models variants. **a** Training loss as a function of training epochs for both model types and tasks, and across BP learning rates. Shaded areas represent s.e.m. across 5 random initializations. **b** Validation loss for a variety of switching model variants for each task, for the best stable BP learning rate found based on **a**. Variants color-highlighted at the top of the panel. Vanilla and switching models from Fig. 2 maintained the same colors. Shaded areas represent s.e.m. across 5 random initializations. **c** Percent BP recurrent weights throughout training for the same model variants as in **b**. Shaded areas represent s.e.m. across 5 random initializations. **d** Flux of synapses switching from Hebb to BP and from BP to Hebb throughout training, for various network sizes for each task. Flux is reported as percentage of total synapses in the network. Shaded areas represent s.e.m. across 5 random initializations. **e** Histogram of the individual synapse Hebbian learning rates Λ*_i,j_* after training, for each BP learning rate, across all network sizes, all tasks, and all random initializations. Histograms are not truncated or otherwise transformed.

**Table S1:** Performance and learning speed for models of size 128. Details are the same as in Tab. 1 (**a**) and Tab. 2 (**b**).

|  |  |  |  |  |  |  |  |  |  |  |
| --- | --- | --- | --- | --- | --- | --- | --- | --- | --- | --- |
| Learning Rate | 1 | 1.5<br>$\pm 0.0$ | 1.5<br>$\pm 0.1$ | 1.3<br>$\pm 0.1$ | 1.3<br>$\pm 0.1$ | 9.1<br>$\pm 0.4$ | 8.3<br>$\pm 0.4$ | 41.7<br>$\pm 1.2$ | 51.3<br>$\pm 2.4$ | 52.0<br>$\pm 2.5$ |
| | | 1.6<br>$\pm 0.1$ | 1.6<br>$\pm 0.1$ | 1.7<br>$\pm 0.1$ | 1.7<br>$\pm 0.0$ | 10.5<br>$\pm 0.7$ | 8.9<br>$\pm 0.6$ | 57.8<br>$\pm 5.5$ | 69.2<br>$\pm 3.0$ | 75.6<br>$\pm 5.8$ |
| | 3.16 | **0.8**<br>**$\pm 0.0$** | **0.8**<br>**$\pm 0.0$** | **0.8**<br>**$\pm 0.1$** | **0.8**<br>**$\pm 0.0$** | 4.7<br>$\pm 0.2$ | 4.6<br>$\pm 0.1$ | **32.2**<br>**$\pm 7.4$** | 35.1<br>$\pm 4.4$ | 45.7<br>$\pm 5.3$ |
| | | 0.8<br>$\pm 0.1$ | 0.9<br>$\pm 0.0$ | 1.0<br>$\pm 0.1$ | 1.0<br>$\pm 0.0$ | 5.1<br>$\pm 0.4$ | 4.8<br>$\pm 0.4$ | 34.2<br>$\pm 5.9$ | **34.5**<br>**$\pm 9.6$** | **39.8**<br>**$\pm 8.6$** |
| | 10 | 1.0<br>$\pm 0.0$ | 1.0<br>$\pm 0.1$ | 1.2<br>$\pm 0.1$ | 1.2<br>$\pm 0.1$ | 4.6<br>$\pm 0.3$ | **4.4**<br>**$\pm 0.2$** | 312.1<br>$\pm 142.1$ | 229.4<br>$\pm 60.9$ | 196.0<br>$\pm 61.7$ |
| | | 0.9<br>$\pm 0.1$ | 1.0<br>$\pm 0.1$ | 1.5<br>$\pm 0.1$ | 1.6<br>$\pm 0.1$ | **4.3**<br>**$\pm 0.2$** | 4.4<br>$\pm 0.1$ | 183.9<br>$\pm 101.8$ | 178.4<br>$\pm 43.7$ | 274.0<br>$\pm 165.8$ |
| | 31.62 | 2.0<br>$\pm 0.2$ | 1.9<br>$\pm 0.2$ | 3.6<br>$\pm 0.8$ | 4.0<br>$\pm 1.2$ | 13.4<br>$\pm 13.2$ | 7.9<br>$\pm 0.6$ | 2121.1<br>$\pm 939.3$ | 2524.0<br>$\pm 210.3$ | 2226.1<br>$\pm 927.1$ |
| | | 1.4<br>$\pm 0.2$ | 1.6<br>$\pm 0.4$ | 4.6<br>$\pm 2.4$ | 9.1<br>$\pm 8.0$ | 1366.7<br>$\pm 1356.5$ | 1756.8<br>$\pm 1343.5$ | 1863.2<br>$\pm 1000.8$ | 2164.3<br>$\pm 1106.8$ | 2636.0<br>$\pm 17.9$ |
|  |  | fdAnti | fdGo | ReactAnti | ReactGo | DelayGo | DelayAnti | MultiDelayDM | DelayDM1 | DelayDM2 |

|  |  |  |  |  |  |  |  |  |  |  |
| --- | --- | --- | --- | --- | --- | --- | --- | --- | --- | --- |
| Learning Rate | 1 | 5.2<br>$\pm 0.3$ | 5.1<br>$\pm 0.2$ | 3.6<br>$\pm 0.1$ | 3.6<br>$\pm 0.2$ | 9.4<br>$\pm 0.6$ | 9.1<br>$\pm 0.8$ | 23.7<br>$\pm 1.9$ | 29.1<br>$\pm 1.2$ | 29.4<br>$\pm 0.8$ |
| | | 6.0<br>$\pm 0.3$ | 6.0<br>$\pm 0.3$ | 3.6<br>$\pm 0.2$ | 3.6<br>$\pm 0.2$ | 10.7<br>$\pm 0.4$ | 10.0<br>$\pm 0.9$ | 34.9<br>$\pm 3.1$ | 43.1<br>$\pm 2.7$ | 42.9<br>$\pm 2.5$ |
| | 3.16 | 2.0<br>$\pm 0.1$ | 2.0<br>$\pm 0.1$ | 1.4<br>$\pm 0.1$ | 1.3<br>$\pm 0.1$ | 3.6<br>$\pm 0.2$ | 3.5<br>$\pm 0.2$ | **11.3**<br>**$\pm 0.9$** | **13.7**<br>**$\pm 0.9$** | **14.1**<br>**$\pm 0.7$** |
| | | 2.3<br>$\pm 0.2$ | 2.3<br>$\pm 0.1$ | 1.3<br>$\pm 0.1$ | 1.3<br>$\pm 0.1$ | 4.2<br>$\pm 0.3$ | 3.9<br>$\pm 0.3$ | 16.0<br>$\pm 1.7$ | 19.2<br>$\pm 0.9$ | 19.3<br>$\pm 1.1$ |
| | 10 | 0.8<br>$\pm 0.0$ | 0.8<br>$\pm 0.0$ | **0.6**<br>**$\pm 0.0$** | **0.6**<br>**$\pm 0.0$** | **1.8**<br>**$\pm 0.1$** | 1.7<br>$\pm 0.1$ | 20.9<br>$\pm 2.2$ | 20.9<br>$\pm 1.5$ | 19.3<br>$\pm 1.4$ |
| | | 0.9<br>$\pm 0.1$ | 0.9<br>$\pm 0.1$ | 0.7<br>$\pm 0.0$ | 0.7<br>$\pm 0.0$ | 2.0<br>$\pm 0.1$ | 2.0<br>$\pm 0.1$ | 18.3<br>$\pm 1.0$ | 19.3<br>$\pm 1.4$ | 19.8<br>$\pm 2.0$ |
| | 31.62 | 0.6<br>$\pm 0.2$ | **0.5**<br>**$\pm 0.0$** | 0.7<br>$\pm 0.1$ | 0.8<br>$\pm 0.2$ | 4.2<br>$\pm 5.7$ | **1.5**<br>**$\pm 0.1$** | 183.5<br>$\pm 82.9$ | 179.8<br>$\pm 76.3$ | 179.9<br>$\pm 79.7$ |
| | | **0.5**<br>**$\pm 0.0$** | 0.5<br>$\pm 0.0$ | 0.9<br>$\pm 0.4$ | 1.2<br>$\pm 0.5$ | 33.8<br>$\pm 51.8$ | 35.2<br>$\pm 35.4$ | 184.0<br>$\pm 56.5$ | 197.7<br>$\pm 83.2$ | 227.9<br>$\pm 14.2$ |
|  |  | fdAnti | fdGo | ReactAnti | ReactGo | DelayGo | DelayAnti | MultiDelayDM | DelayDM1 | DelayDM2 |

**Table S2:**
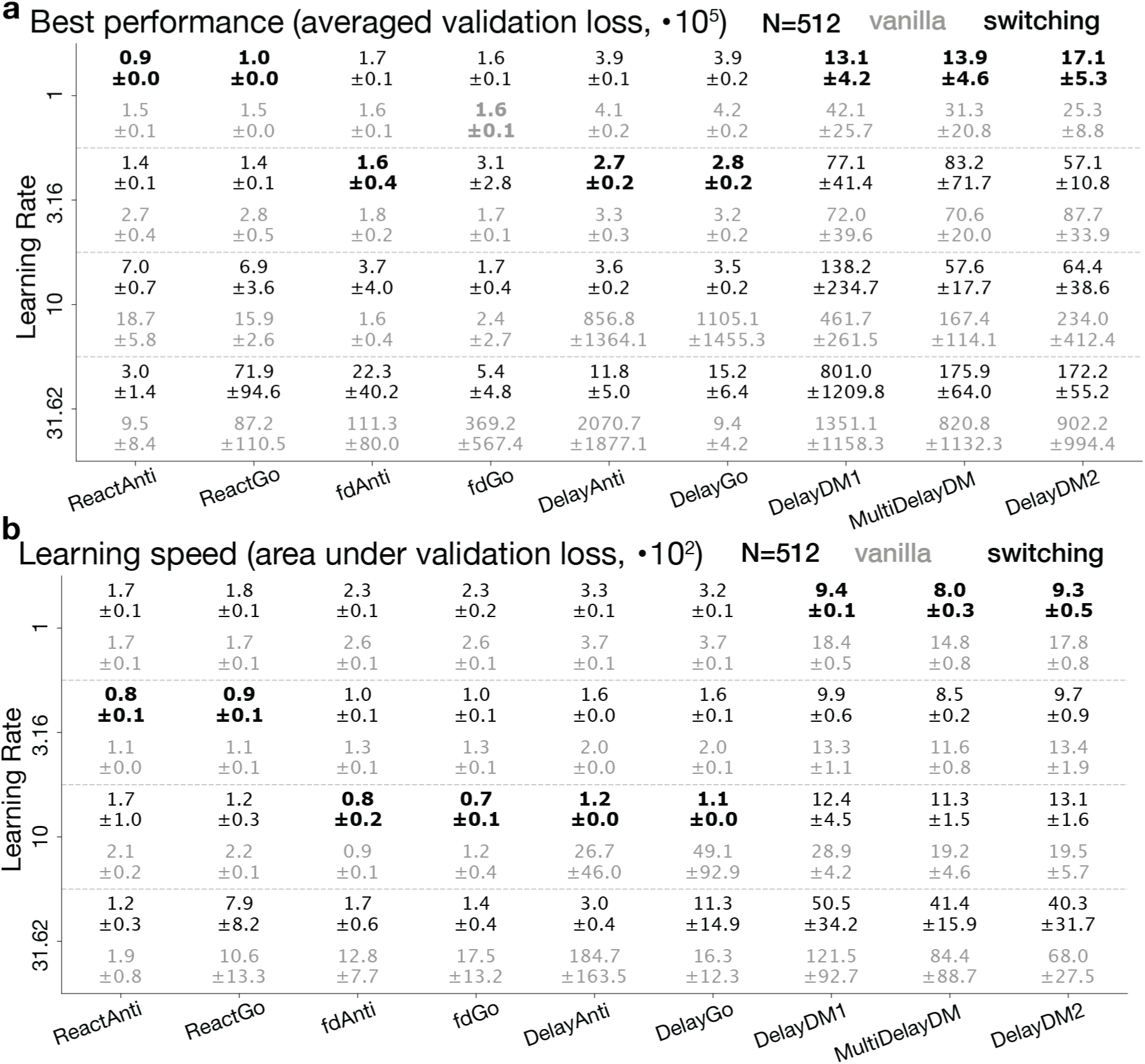
Performance and learning speed for models of size 512. Details are the same as in Tab. 1 (**a**) and Tab. 2 (**b**).

**Table S3:**
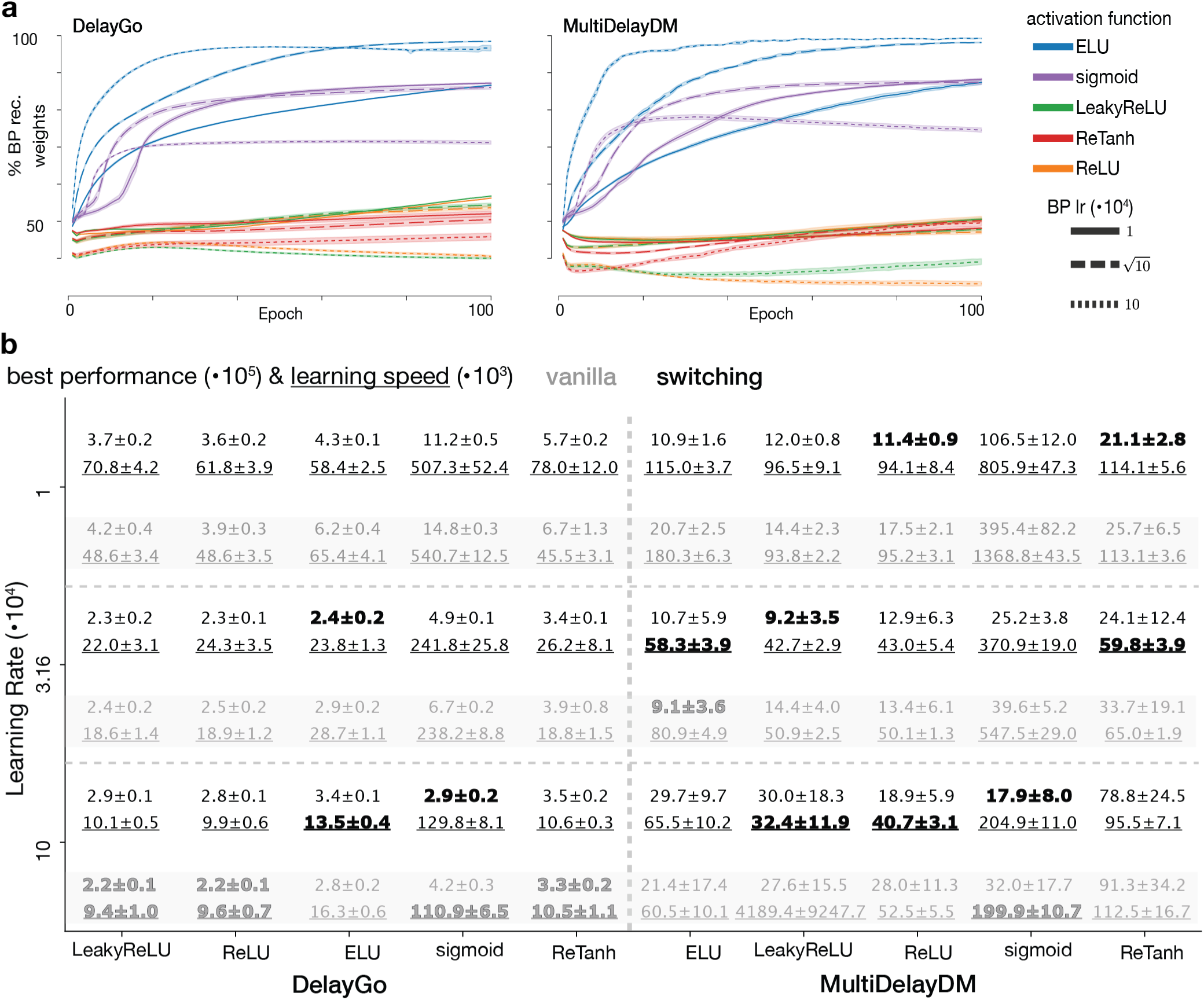
Switching models with various transfer functions. **a** Percent BP recurrent weights throughout training for switching models with different transfer functions (color-coded), across BP learning rates (solid, dashed, dotted lines) for each task (left, right). Shaded areas represent s.e.m. across 5 random initializations. **b** Table summarizing the best performance (as in Tab. 1) and learning speed (as in Tab. 2) for switching models with different transfer functions across BP learning rates for each task. Best performance and learning speed are highlighted in bold for each transfer function and task. Transfer functions are ordered by increasing performance, i.e. best performance across our trained models, from left to right.

**Figure S3:**
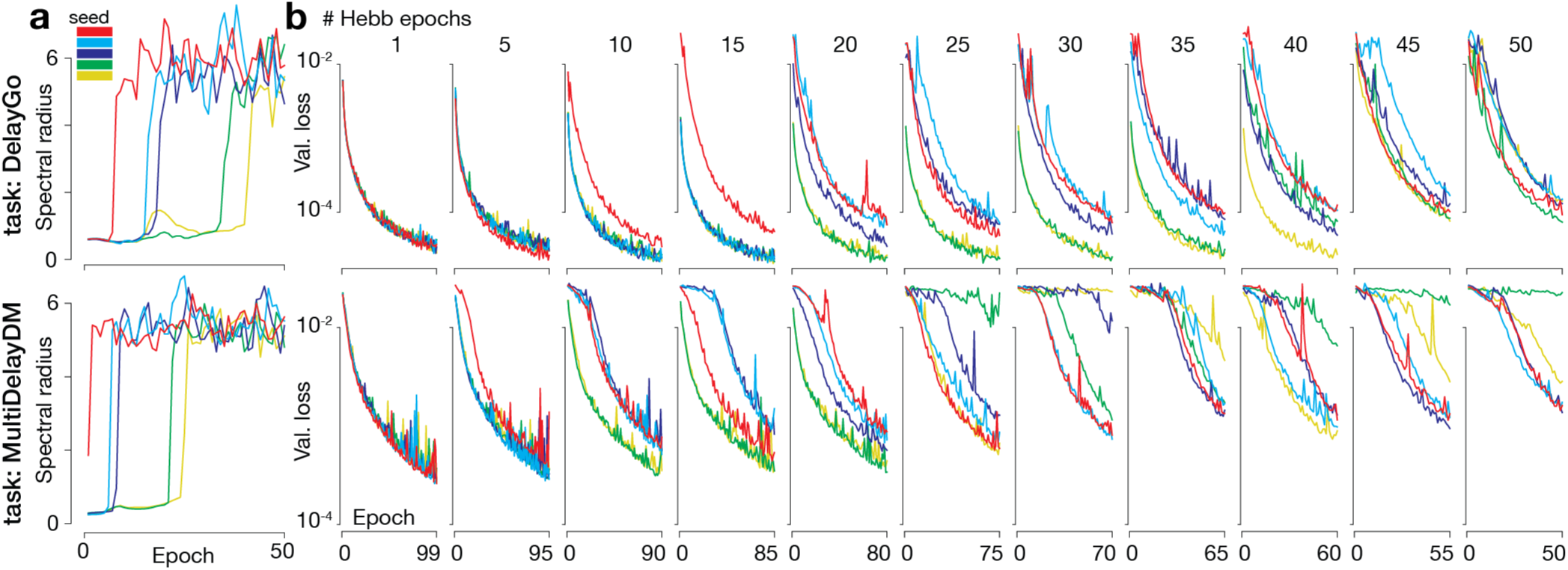
Hebb-only pre-training and its effects on vanilla models. **a** Spectral radius of the recurrent weights matrix throughout training for Hebb-only pre-trained models for each task. 5 individual models are shown in different colors for each random seed. **b** Validation loss throughout training for vanilla models initialized from pre-trained Hebb-only models, color-coded by the same random seed colors as in **a**. Each column corresponds to increasing number of pre-trained epochs of the Hebb-only model used for initialization (top of column, left to right).

**Figure S4:**
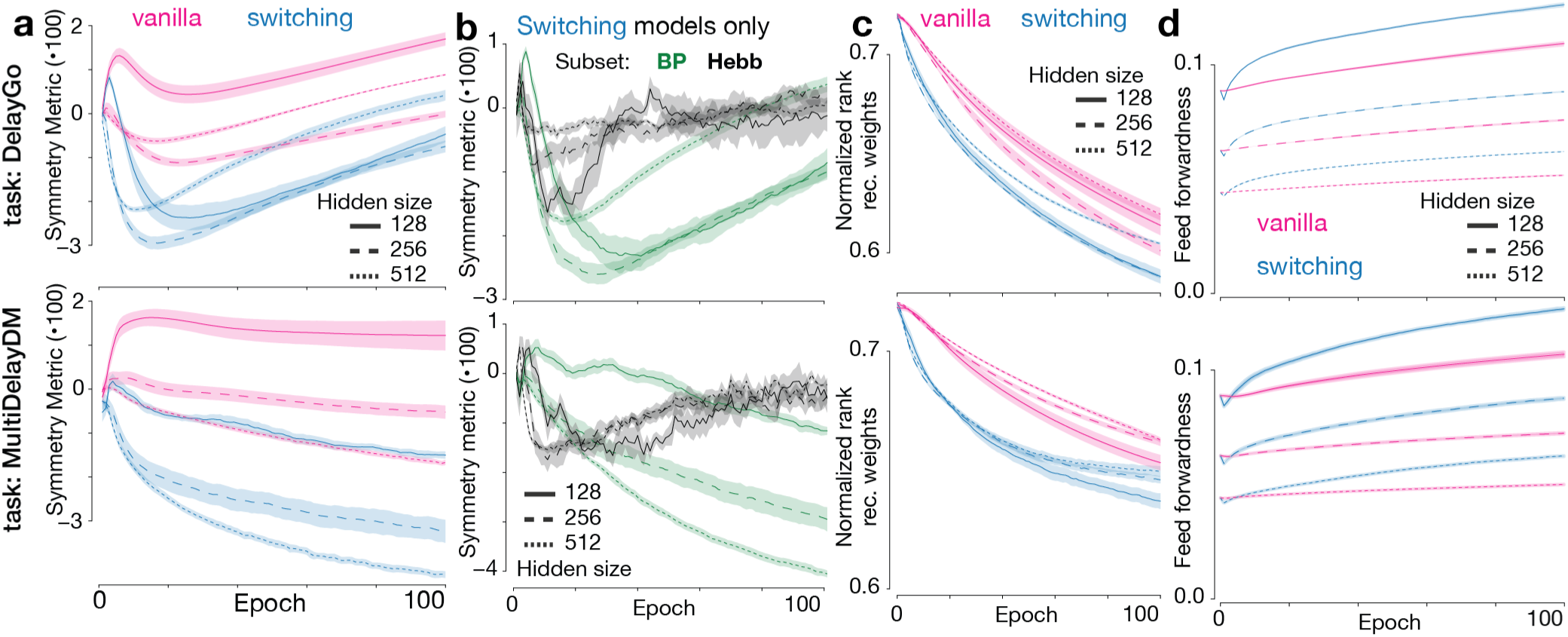
Structural properties across network sizes. Same as Fig. 4, but for different network sizes (solid, dashed, and dotted lines for sizes 128, 256, and 512, respectively) for each task. **a** Symmetry metric of the recurrent weights matrix, **b** symmetry metric of the Hebb and BP subsets of the recurrent weights matrix for switching models, **c** normalized rank of the recurrent weights matrix, and **d** amount of feed forward structure in the recurrent weights matrix, as a function of training epochs for both model types and tasks.

## Notes

https://github.com/DenisTurcu/psRNN

### Competing Interest Statement

The authors have declared no competing interest.

### Summary of Updates

We better explained the contributions for the broader neuroscience community.

## References

[1] J Spacek. Three-dimensional analysis of dendritic spines. ii. spine apparatus and other cytoplasmic components. Anatomy and embryology, 171(2):235–243, 1985.

[2] Thomas Deller, Martin Korte, Sophie Chabanis, Alexander Drakew, Herbert Schwegler, Giulia Good Stefani, Aimee Zuniga, Karin Schwarz, Tobias Bonhoeffer, Rolf Zeller, et al. Synaptopodin-deficient mice lack a spine apparatus and show deficits in synaptic plasticity. Proceedings of the National Academy of Sciences, 100(18):10494–10499, 2003.

[3] Peter Jedlicka, Andreas Vlachos, Stephan W Schwarzacher, and Thomas Deller. A role for the spine apparatus in ltp and spatial learning. Behavioural brain research, 192(1):12–19, 2008.

[4] Niklaus Holbro, Åsa Grunditz, and Thomas G Oertner. Differential distribution of endo-plasmic reticulum controls metabotropic signaling and plasticity at hippocampal synapses. Proceedings of the National Academy of Sciences, 106(35):15055–15060, 2009.

[5] Eugene M Izhikevich and Gerald M Edelman. Large-scale model of mammalian thalamocortical systems. Proceedings of the national academy of sciences, 105(9):3593–3598, 2008.

[6] Tobias C Potjans and Markus Diesmann. The cell-type specific cortical microcircuit: relating structure and activity in a full-scale spiking network model. Cerebral cortex, 24(3):785–806, 2014.

[7] Srikanth Ramaswamy, Jean-Denis Courcol, Marwan Abdellah, Stanislaw R Adaszewski, Nicolas Antille, Selim Arsever, Guy Atenekeng, Ahmet Bilgili, Yury Brukau, Athanassia Chalimourda, et al. The neocortical microcircuit collaboration portal: a resource for rat somatosensory cortex. Frontiers in neural circuits, 9:44, 2015.

[8] Loreen Hertäg and Henning Sprekeler. Amplifying the redistribution of somato-dendritic inhibition by the interplay of three interneuron types. PLoS computational biology, 15(5):e1006999, 2019.

[9] Matt Udakis, Victor Pedrosa, Sophie EL Chamberlain, Claudia Clopath, and Jack R Mellor. Interneuron-specific plasticity at parvalbumin and somatostatin inhibitory synapses onto ca1 pyramidal neurons shapes hippocampal output. Nature communications, 11(1):4395, 2020.

[10] Sean Froudist-Walsh, Daniel P Bliss, Xingyu Ding, Lucija Rapan, Meiqi Niu, Kenneth Knoblauch, Karl Zilles, Henry Kennedy, Nicola Palomero-Gallagher, and Xiao-Jing Wang. A dopamine gradient controls access to distributed working memory in the large-scale monkey cortex. Neuron, 109(21):3500–3520, 2021.

[11] Liqun Luo. Architectures of neuronal circuits. Science, 373(6559):eabg7285, 2021.

[12] Gord Fishell and Adam Kepecs. Interneuron types as attractors and controllers. Annual review of neuroscience, 43(1):1–30, 2020.

[13] Sven Dorkenwald, Nicholas L Turner, Thomas Macrina, Kisuk Lee, Ran Lu, Jingpeng Wu, Agnes L Bodor, Adam A Bleckert, Derrick Brittain, Nico Kemnitz, et al. Binary and analog variation of synapses between cortical pyramidal neurons. Elife, 11:e76120, 2022.

[14] Henry Markram, Joachim Lübke, Michael Frotscher, and Bert Sakmann. Regulation of synaptic efficacy by coincidence of postsynaptic aps and epsps. Science, 275(5297):213–215, 1997.

[15] Guo-qiang Bi and Muming Poo. Synaptic modifications in cultured hippocampal neurons: dependence on spike timing, synaptic strength, and postsynaptic cell type. Journal of neuroscience, 18(24):10464–10472, 1998.

[16] Guo-qiang Bi and Mu-ming Poo. Synaptic modification by correlated activity: Hebb’s postulate revisited. Annual review of neuroscience, 24(1):139–166, 2001.

[17] Natalia Caporale and Yang Dan. Spike timing–dependent plasticity: a hebbian learning rule. Annu. Rev. Neurosci., 31(1):25–46, 2008.

[18] Wulfram Gerstner. Hebbian learning and plasticity. From neuron to cognition via computational neuroscience, pages 0–25, 2011.

[19] Henry Markram, Wulfram Gerstner, and Per Jesper Sjöström. A history of spike-timing-dependent plasticity. Frontiers in synaptic neuroscience, 3:4, 2011.

[20] Wulfram Gerstner, Richard Kempter, J Leo Van Hemmen, and Hermann Wagner. A neuronal learning rule for sub-millisecond temporal coding. Nature, 383(6595):76–78, 1996.

[21] Sen Song, Kenneth D Miller, and Larry F Abbott. Competitive hebbian learning through spike-timing-dependent synaptic plasticity. Nature neuroscience, 3(9):919–926, 2000.

[22] Tim P Vogels, Henning Sprekeler, Friedemann Zenke, Claudia Clopath, and Wulfram Gerstner. Inhibitory plasticity balances excitation and inhibition in sensory pathways and memory networks. Science, 334(6062):1569–1573, 2011.

[23] James A D’amour and Robert C Froemke. Inhibitory and excitatory spike-timing-dependent plasticity in the auditory cortex. Neuron, 86(2):514–528, 2015.

[24] Fereshteh Lagzi, Martha Canto Bustos, Anne-Marie Oswald, and Brent Doiron. Assembly formation is stabilized by parvalbumin neurons and accelerated by somatostatin neurons. BioRxiv, pages 2021–09, 2021.

[25] Basile Confavreux, Everton J Agnes, Friedemann Zenke, Henning Sprekeler, and Tim P Vogels. Balancing complexity, performance and plausibility to meta learn plasticity rules in recurrent spiking networks. PLOS Computational Biology, 21(4):e1012910, 2025.

[26] Francis Crick. The recent excitement about neural networks. Nature, 337(6203):129–132, 1989.

[27] James L McClelland. How far can you go with hebbian learning, and when does it lead you astray. Processes of change in brain and cognitive development: Attention and performance xxi, 21:33–69, 2006.

[28] Yoshua Bengio, Dong-Hyun Lee, Jorg Bornschein, Thomas Mesnard, and Zhouhan Lin. Towards biologically plausible deep learning. *arXiv preprint arXiv:1502.04156*, 2015.

[29] Nicolas Frémaux and Wulfram Gerstner. Neuromodulated spike-timing-dependent plasticity, and theory of three-factor learning rules. Frontiers in neural circuits, 9:85, 2016.

[30] Pieter R Roelfsema and Anthony Holtmaat. Control of synaptic plasticity in deep cortical networks. Nature Reviews Neuroscience, 19(3):166–180, 2018.

[31] Yicong Zheng, Xiaonan L Liu, Satoru Nishiyama, Charan Ranganath, and Randall C O’Reilly. Correcting the hebbian mistake: Toward a fully error-driven hippocampus. PLOS Computational Biology, 18(10):e1010589, 2022.

[32] Timothy P Lillicrap, Daniel Cownden, Douglas B Tweed, and Colin J Akerman. Random synaptic feedback weights support error backpropagation for deep learning. Nature communications, 7(1):13276, 2016.

[33] João Sacramento, Rui Ponte Costa, Yoshua Bengio, and Walter Senn. Dendritic cortical microcircuits approximate the backpropagation algorithm. Advances in neural information processing systems, 31, 2018.

[34] Blake A Richards and Timothy P Lillicrap. Dendritic solutions to the credit assignment problem. Current opinion in neurobiology, 54:28–36, 2019.

[35] Timothy P Lillicrap, Adam Santoro, Luke Marris, Colin J Akerman, and Geoffrey Hinton. Backpropagation and the brain. Nature Reviews Neuroscience, 21(6):335–346, 2020.

[36] Łukasz Kuśmierz, Takuya Isomura, and Taro Toyoizumi. Learning with three factors: modulating hebbian plasticity with errors. Current opinion in neurobiology, 46:170–177, 2017.

[37] Sergey Bartunov, Adam Santoro, Blake Richards, Luke Marris, Geoffrey E Hinton, and Timothy Lillicrap. Assessing the scalability of biologically-motivated deep learning algorithms and architectures. Advances in neural information processing systems, 31, 2018.

[38] Stefano Fusi, Patrick J Drew, and Larry F Abbott. Cascade models of synaptically stored memories. Neuron, 45(4):599–611, 2005.

[39] Danil Tyulmankov, Guangyu Robert Yang, and LF Abbott. Meta-learning synaptic plasticity and memory addressing for continual familiarity detection. Neuron, 110(3):544–557, 2022.

[40] Kyle Aitken and Stefan Mihalas. Neural population dynamics of computing with synaptic modulations. Elife, 12:e83035, 2023.

[41] James CR Whittington, Timothy H Muller, Shirley Mark, Guifen Chen, Caswell Barry, Neil Burgess, and Timothy EJ Behrens. The tolman-eichenbaum machine: unifying space and relational memory through generalization in the hippocampal formation. Cell, 183(5):1249–1263, 2020.

[42] Roman Pogodin, Jonathan Cornford, Arna Ghosh, Gauthier Gidel, Guillaume Lajoie, and Blake Richards. Synaptic weight distributions depend on the geometry of plasticity. *arXiv preprint arXiv:2305.19394*, 2023.

[43] Jonathan Cornford, Roman Pogodin, Arna Ghosh, Kaiwen Sheng, Brendan A Bicknell, Olivier Codol, Beverley A Clark, Guillaume Lajoie, and Blake A Richards. Brain-like learning with exponentiated gradients. bioRxiv, pages 2024–10, 2024.

[44] Yann LeCun, Léon Bottou, Genevieve B Orr, and Klaus-Robert Müller. Efficient backprop. In Neural networks: Tricks of the trade, pages 9–50. Springer, 2002.

[45] Xavier Glorot and Yoshua Bengio. Understanding the difficulty of training deep feedforward neural networks. In Proceedings of the thirteenth international conference on artificial intelligence and statistics, pages 249–256. JMLR Workshop and Conference Proceedings, 2010.

[46] Ilya Sutskever, James Martens, George Dahl, and Geoffrey Hinton. On the importance of initialization and momentum in deep learning. In International conference on machine learning, pages 1139–1147. pmlr, 2013.

[47] Dmytro Mishkin and Jiri Matas. All you need is a good init. *arXiv preprint arXiv:1511.06422*, 2015.

[48] Hongyi Zhang, Yann N Dauphin, and Tengyu Ma. Fixup initialization: Residual learning without normalization. *arXiv preprint arXiv:1901.09321*, 2019.

[49] Guangyu Robert Yang, Madhura R Joglekar, H Francis Song, William T Newsome, and Xiao-Jing Wang. Task representations in neural networks trained to perform many cognitive tasks. Nature neuroscience, 22(2):297–306, 2019.

[50] Lea Duncker, Laura Driscoll, Krishna V Shenoy, Maneesh Sahani, and David Sussillo. Organizing recurrent network dynamics by task-computation to enable continual learning. Advances in neural information processing systems, 33:14387–14397, 2020.

[51] Ta-Chu Kao, Kristopher Jensen, Gido Van De Ven, Alberto Bernacchia, and Guillaume Hennequin. Natural continual learning: success is a journey, not (just) a destination. Advances in neural information processing systems, 34:28067–28079, 2021.

[52] Mikail Khona, Sarthak Chandra, Joy J Ma, and Ila R Fiete. Winning the lottery with neural connectivity constraints: Faster learning across cognitive tasks with spatially constrained sparse rnns. Neural Computation, 35(11):1850–1869, 2023.

[53] Laura N Driscoll, Krishna Shenoy, and David Sussillo. Flexible multitask computation in recurrent networks utilizes shared dynamical motifs. Nature Neuroscience, 27(7):1349–1363, 2024.

[54] Haozhe Shan, Sun Minni, and Lea Duncker. Separating the what and how of compositional computation to enable reuse and continual learning. arXiv preprint arXiv:2510.20709, 2025.

[55] Shintaro Funahashi, Charles J Bruce, and Patricia S Goldman-Rakic. Mnemonic coding of visual space in the monkey’s dorsolateral prefrontal cortex. Journal of neurophysiology, 61(2):331–349, 1989.

[56] David Raposo, Matthew T Kaufman, and Anne K Churchland. A category-free neural population supports evolving demands during decision-making. Nature neuroscience, 17(12):1784–1792, 2014.

[57] Robert C Froemke, Mu-ming Poo, and Yang Dan. Spike-timing-dependent synaptic plasticity depends on dendritic location. Nature, 434(7030):221–225, 2005.

[58] Johannes J Letzkus, Bjö rn M Kampa, and Greg J Stuart. Learning rules for spike timing-dependent plasticity depend on dendritic synapse location. Journal of Neuroscience, 26(41):10420–10429, 2006.

[59] Robert C Froemke, Johannes J Letzkus, Bjö rn M Kampa, Giao B Hang, and Greg J Stuart. Dendritic synapse location and neocortical spike-timing-dependent plasticity. Frontiers in synaptic neuroscience, 2:29, 2010.

[60] Aparna Suvrathan. Beyond stdp—towards diverse and functionally relevant plasticity rules. Current opinion in neurobiology, 54:12–19, 2019.

[61] Aref Pariz, Daniel Trotter, Axel Hutt, and Jeremie Lefebvre. Selective control of synaptic plasticity in heterogeneous networks through transcranial alternating current stimulation (tacs). PLOS Computational Biology, 19(4):e1010736, 2023.

[62] Mantas Lukoševičius and Herbert Jaeger. Reservoir computing approaches to recurrent neural network training. Computer science review, 3(3):127–149, 2009.

[63] Mantas Lukoševičius, Herbert Jaeger, and Benjamin Schrauwen. Reservoir computing trends. KI-Kü nstliche Intelligenz, 26(4):365–371, 2012.

[64] Gouhei Tanaka, Toshiyuki Yamane, Jean Benoit Héroux, Ryosho Nakane, Naoki Kanazawa, Seiji Takeda, Hidetoshi Numata, Daiju Nakano, and Akira Hirose. Recent advances in physical reservoir computing: A review. Neural Networks, 115:100–123, 2019.

[65] Brian Hu, Marina E Garrett, Peter A Groblewski, Douglas R Ollerenshaw, Jiaqi Shang, Kate Roll, Sahar Manavi, Christof Koch, Shawn R Olsen, and Stefan Mihalas. Adaptation supports short-term memory in a visual change detection task. PLoS computational biology, 17(9):e1009246, 2021.

[66] Marcelo O Magnasco, Oreste Piro, and Guillermo A Cecchi. Self-tuned critical anti-hebbian networks. Physical review letters, 102(25):258102, 2009.

[67] Lee Susman, Naama Brenner, and Omri Barak. Stable memory with unstable synapses. Nature communications, 10(1):4441, 2019.

[68] Joel T Kaardal, Frédéric E Theunissen, and Tatyana O Sharpee. A low-rank method for characterizing high-level neural computations. Frontiers in computational neuroscience, 11:68, 2017.

[69] Francesca Mastrogiuseppe and Srdjan Ostojic. Linking connectivity, dynamics, and computations in low-rank recurrent neural networks. Neuron, 99(3):609–623, 2018.

[70] Friedrich Schuessler, Francesca Mastrogiuseppe, Alexis Dubreuil, Srdjan Ostojic, and Omri Barak. The interplay between randomness and structure during learning in rnns. Advances in neural information processing systems, 33:13352–13362, 2020.

[71] Alexis Dubreuil, Adrian Valente, Manuel Beiran, Francesca Mastrogiuseppe, and Srdjan Ostojic. The role of population structure in computations through neural dynamics. Nature neuroscience, 25(6):783–794, 2022.

[72] Edward J Hu, Yelong Shen, Phillip Wallis, Zeyuan Allen-Zhu, Yuanzhi Li, Shean Wang, Liang Wang, Weizhu Chen, et al. Lora: Low-rank adaptation of large language models. Iclr, 1(2):3, 2022.

[73] Ruixiang Zhang, Shuangfei Zhai, Etai Littwin, and Josh Susskind. Learning representation from neural fisher kernel with low-rank approximation. *arXiv preprint arXiv:2202.01944*, 2022.

[74] Arthur Pellegrino, N Alex Cayco Gajic, and Angus Chadwick. Low tensor rank learning of neural dynamics. Advances in Neural Information Processing Systems, 36:11674–11702, 2023.

[75] Manuel Beiran, Nicolas Meirhaeghe, Hansem Sohn, Mehrdad Jazayeri, and Srdjan Ostojic. Parametric control of flexible timing through low-dimensional neural manifolds. Neuron, 111(5):739–753, 2023.

[76] Samuel Horváth, Stefanos Laskaridis, Shashank Rajput, and Hongyi Wang. Maestro: Uncovering low-rank structures via trainable decomposition. *arXiv preprint arXiv:2308.14929*, 2023.

[77] Francesca Mastrogiuseppe, Joana Carmona, and Christian K Machens. Stochastic activity in low-rank recurrent neural networks. PLOS Computational Biology, 21(8):e1013371, 2025.

[78] Jia-Chen Zhang, Yu-Jie Xiong, Chun-Ming Xia, Dong-Hai Zhu, and Hong-Jian Zhan. Lora2: Multi-scale low-rank approximations for fine-tuning large language models. Neurocomputing, 650:130859, 2025.

[79] Laura Balzano, Tianjiao Ding, Benjamin D Haeffele, Soo Min Kwon, Qing Qu, Peng Wang, Zhangyang Wang, and Can Yaras. An overview of low-rank structures in the training and adaptation of large models. *arXiv preprint arXiv:2503.19859*, 2025.

[80] Vladimir K Berezovskii, Jonathan J Nassi, and Richard T Born. Segregation of feedforward and feedback projections in mouse visual cortex. Journal of Comparative Neurology, 519(18):3672–3683, 2011.

[81] Boris Vladimirskiy, Robert Urbanczik, and Walter Senn. Hierarchical novelty-familiarity representation in the visual system by modular predictive coding. PLoS One, 10(12):e0144636, 2015.

[82] Jorge F Mejias, John D Murray, Henry Kennedy, and Xiao-Jing Wang. Feedforward and feedback frequency-dependent interactions in a large-scale laminar network of the primate cortex. Science advances, 2(11):e1601335, 2016.

[83] Ruben S van Bergen and Nikolaus Kriegeskorte. Going in circles is the way forward: the role of recurrence in visual inference. Current Opinion in Neurobiology, 65:176–193, 2020.

[84] Hirokazu Tanaka, Takahiro Ishikawa, Jongho Lee, and Shinji Kakei. The cerebro-cerebellum as a locus of forward model: a review. Frontiers in systems neuroscience, 14:19, 2020.

[85] Joseph Pemberton, Ellen Boven, Richard Apps, and Rui Ponte Costa. Cortico-cerebellar networks as decoupling neural interfaces. Advances in neural information processing systems, 34:7745–7759, 2021.

[86] Elham Barzegaran and Gijs Plomp. Multiple concurrent feedforward and feedback streams in a cortical hierarchy. bioRxiv, pages 2021–01, 2021.

[87] Ran Wang, Xupeng Chen, Amirhossein Khalilian-Gourtani, Leyao Yu, Patricia Dugan, Daniel Friedman, Werner Doyle, Orrin Devinsky, Yao Wang, and Adeen Flinker. Distributed feedforward and feedback cortical processing supports human speech production. Proceedings of the National Academy of Sciences, 120(42):e2300255120, 2023.

[88] Siming Yan, Xuyang Fang, Bowen Xiao, Harold Rockwell, Yimeng Zhang, and Tai Sing Lee. Recurrent feedback improves feedforward representations in deep neural networks. *arXiv preprint arXiv:1912.10489*, 2019.

[89] Martin Do Pham, Amedeo D’Angiulli, Maryam Mehri Dehnavi, and Robin Chhabra. From brain models to robotic embodied cognition: How does biological plausibility inform neuro-morphic systems? Brain Sciences, 13(9):1316, 2023.

[90] Felix A Wichmann and Robert Geirhos. Are deep neural networks adequate behavioral models of human visual perception? Annual review of vision science, 9(1):501–524, 2023.

[91] Peter Henrici. Bounds for iterates, inverses, spectral variation and fields of values of non-normal matrices. Numerische Mathematik, 4(1):24–40, 1962.

[92] Neta Ravid Tannenbaum and Yoram Burak. Shaping neural circuits by high order synaptic interactions. PLOS Computational Biology, 12(8):e1005056, 2016.

[93] James A Henderson and Pulin Gong. Functional mechanisms underlie the emergence of a diverse range of plasticity phenomena. PLoS Computational Biology, 14(11):e1006590, 2018.

[94] John J Hopfield. Neural networks and physical systems with emergent collective computational abilities. Proceedings of the national academy of sciences, 79(8):2554–2558, 1982.

[95] Jinwen Ma. The asymptotic memory capacity of the generalized hopfield network. Neural Networks, 12(9):1207–1212, 1999.

[96] Diederik P Kingma and Jimmy Ba. Adam: A method for stochastic optimization. *arXiv preprint arXiv:1412*.6980, 2014.

[97] Malbor Asllani, Renaud Lambiotte, and Timoteo Carletti. Structure and dynamical behavior of non-normal networks. Science advances, 4(12):eaau9403, 2018.

